# Hypoxia adaptation shapes genomic architecture and vertical niche transitions in copepods

**DOI:** 10.1101/2025.11.05.686523

**Authors:** Kévin Sugier, Romuald Laso-Jadart, Loïc Dorval, Arnaud Meng, Rainer Kiko, Leocadio Blanco-Bercial, Amy E. Maas, Astrid Cornils, Frédéric Maps, Sakina-Dorothée Ayata, Mohammed-Amin Madoui

## Abstract

Oxygen Minimum Zone (OMZ) expansion is a major challenge to marine ecosystems and associated zooplankton. Calanoid copepods include lineages that tolerate hypoxia and exhibit functional traits such as diel vertical migrations towards, or dormancy within, hypoxic mesopelagic zones. However, the evolutionary origins and molecular drivers of these functional traits remain unclear. Herein, we integrate a time-calibrated phylotranscriptomic tree of 50 copepod species with ancestral trait reconstruction, gene family copy number variation, and palaeoceanographic data to infer the evolutionary timing and ecological drivers of hypoxia adaptation. Our results support the following scenario: 1. Post-embryonic dormancy probably originated in calanoid ancestors, accompanied by widespread gene expansions involving hypoxia-response pathways as well as lipid and amino acid metabolisms. 2. Mesopelagic colonisation by calanoid lineages likely occurred during the Ordovician deep-sea oxygenation event. 3. During the Carboniferous deep-sea deoxygenation, a secondary habitat shift would have occurred toward shallower waters and embryonic dormancy, and as well as a contraction of the size of gene families in the Diaptomoidae. We further analysed the transcriptomic response of *Eucalanus hyalinus* from the Benguela upwelling OMZ versus surface waters, and identified a coordinated response involving extracellular matrix remodelling, amino acid recycling for anaerobic energy and antioxidant production as well as triglycerides to wax ester conversion. Ka/Ks analysis of gene family expansions upstream (proteolysis, transport) and downstream (antioxidant biosynthesis) of core metabolic pathways suggested purifying selection on dosage-sensitive genes. Together, these results link palaeoclimate change to lineage-specific genome evolution patterns supporting copepod adaptation to oxygen limitation.

## Introduction

Oxygen Minimum Zones (OMZs) are midwater ocean layers (∼100–1,000 m) where dissolved oxygen concentrations reach very low levels due to limited ventilation and high microbial respiration of sinking organic matter^1,2^. OMZs are currently expanding and represent a major threat to marine biodiversity and ecosystem stability, particularly in pelagic communities that are vulnerable to declining oxygen availability^3,4^. Zooplankton taxa that tolerate hypoxia, such as certain calanoid copepods, play a critical role in marine food webs and in regulating global biogeochemical cycles through carbon sequestration and nutrient cycling^5^. These copepods often dominate the zooplankton biomass and certain taxa exhibit a suite of functional traits, like embryonic or post-embryonic dormancy and diel vertical migration (DVM) into the mesopelagic zone (200 to 2,000 m), that facilitate adaptation to low-oxygen environments and may confer resilience to OMZ expansion^6,7^.

Dormancy in copepods has been extensively documented as a key adaptive strategy to cope with environmental variability. It encompasses a spectrum of reversible developmental arrests, including embryonic dormancy (resting eggs delaying hatching), post-embryonic dormancy (occurring after hatching), and diapause (a recurrent, genetically programmed and long-term post-embryonic dormancy that occurs in adults and at late juvenile stages)^8,9^. DVM is characterised by daily vertical migration across depth gradients. Usually, individuals ascend to surface waters at night to feed while remaining hidden from visual predators, and descend to deeper layers during the day to reduce predation risk. In copepods, DVM can span tens to hundreds of meters and exposes organisms to strong environmental gradients, often involving regular excursions into OMZs^10^.

During both dormancy and diel vertical migration (DVM), copepods encounter hypoxia, defined as dissolved oxygen concentrations below ∼2 mg O_2_ L^−1^ in marine systems^10,11^. Hypoxia tolerance corresponds to the capacity to withstand short-term oxygen limitation while minimising cellular damage and is widely considered an ancestral feature of metazoans, consistent with early evolution under low-oxygen conditions^12^. At the molecular level, this response is primarily mediated by the Hypoxia-inducible factor 1 (HIF-1) pathway^13^, a conserved regulator of oxygen sensing across protostomes, vertebrates, and cnidarians, controlling genes involved in anaerobic metabolism, angiogenesis, and cellular survival^14^. Notably, HIF-1 has been secondarily lost in non-diapausing copepods like harpacticoids and cyclopoids, while it is retained in calanoids^15,16^. Gene duplication within hypoxia-responsive pathways has been demonstrated to increase hypoxia tolerance through gene dosage effects, subfunctionalisation, or neofunctionalisation, leading to an by extension of the duration or an increase in intensity of hypoxia resistance in cyprinid fishes^17^ and molluscs^18^. In contrast to hypoxia tolerance, hypoxia adaptation refers to heritable, lineage-specific modifications that enhance performance under chronic or recurrent hypoxia^19^.

To better understand present-day adaptation to hypoxia in copepods, it is necessary to consider how oceanic palaeoclimate changes have shaped their evolutionary history. Two decades ago, Bradford-Grieve^20^ proposed that calanoid copepods colonised the pelagic realm in successive invasions associated with major paleoceanographic transitions, with Arietelloidea and Diaptomoidea invading during the Devonian, and Calanoidea–Clausocalanoidea arising in the Permian during deep-ocean ventilation. Ten years later, Blanco-Bercial and colleagues^21^ reconstructed a molecular phylogeny based on two nuclear and two mitochondrial markers, confirming the monophyly of Calanoida and resolving most superfamilies as coherent lineages, but without explicit temporal staging of mesopelagic invasions. These works were based on oceanic paleoclimate models, which were recently improved for reconstruction of deep-sea oxygen concentration^22^. Also, the tree topology inferred by Blanco-Bercial et *al*. was based on few markers that did not resolve deep nodes.

In the present study, we tested the Bradford-Grieve hypothesis by inferring the evolutionary timing and ecological drivers of hypoxia adaptation and associated gene family expansion in copepods. We integrated a time-calibrated phylotranscriptomic tree of 50 copepod species, including six newly sequenced taxa, with ancestral traits reconstruction, gene family copy number variation, and oceanic paleoclimate data. We further conducted differential gene expression analysis and manual functional annotation of *Eucalanus hyalinus* under *in situ* hypoxic conditions, allowing the identification of specific genes and physiological processes associated with hypoxia tolerance, as well as the evolutionary dynamics of hypoxia response gene paralogues across calanoid lineages. This integrated approach provides a mechanistic and macroevolutionary understanding of copepod resilience to hypoxia and the origins of key functional traits in marine zooplankton.

## Results and discussion

### Phylotranscriptomic tree and trait evolution in copepods

We assembled transcriptomes for 50 copepod species, including six newly sequenced taxa and 44 species from published RNA-seq datasets (Supplementary Table 1). The phylotranscriptomic tree inferred from a concatenated alignment of 1,322 orthologous genes strongly supports the emergence of Metridinidae prior to the divergence of Pancalanina (a provisional suborder level group proposed here to encompass Calanidae, Clausocalanidae, Eucalanidae, Euchaetidae, and Rhincalanidae, and their corresponding superfamilies) and Diapotomoidea (bootstrap support = 100; Fig. 1a). This phylotranscriptomic tree provides a robust topology similar and more complete than previous reconstructions^23^ and enables confident reconstruction of ancestral hypoxia-associated functional traits (Supplementary Tables 2,3).

**Fig. 1.**
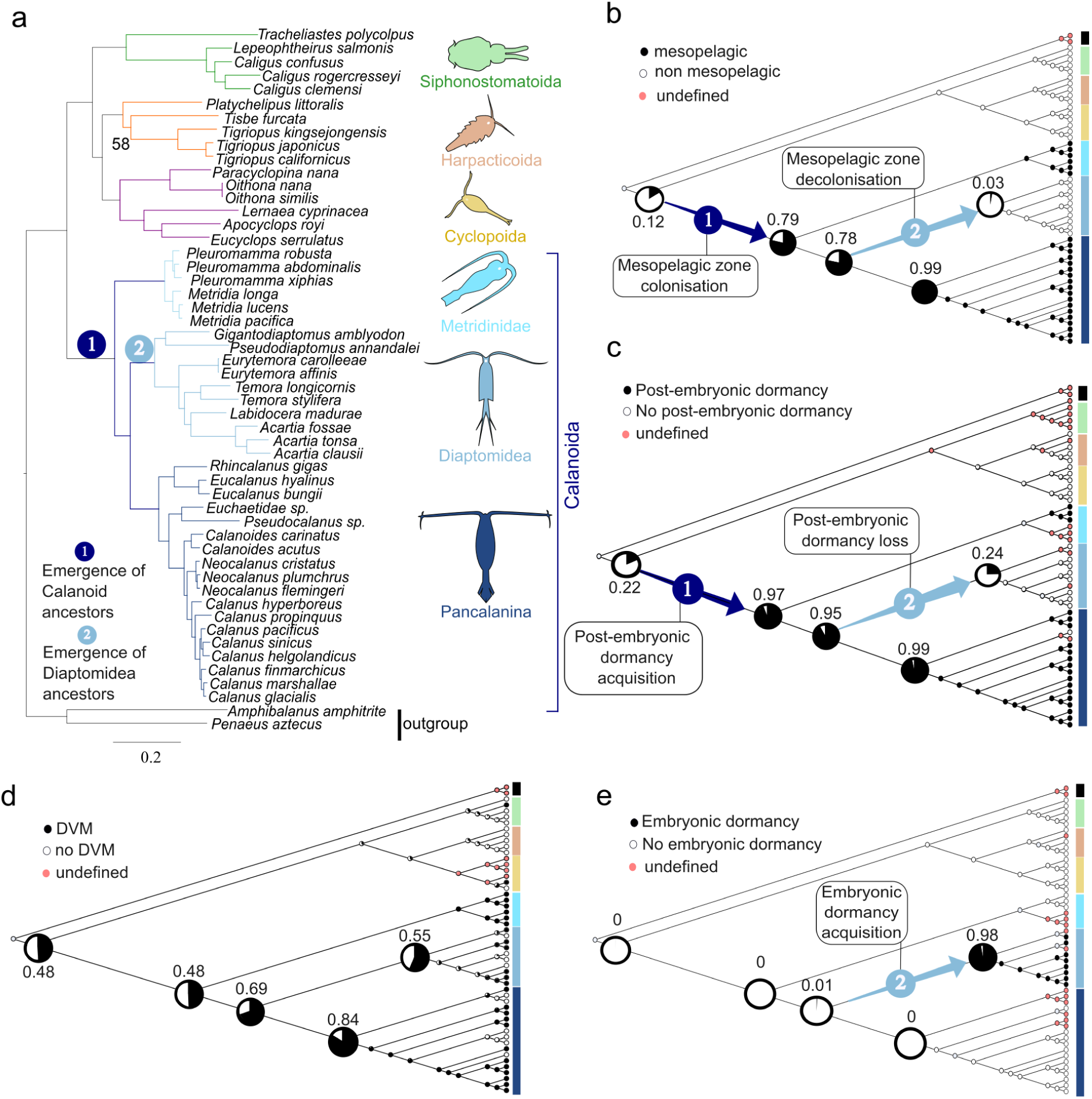
Molecular and trait evolution of copepods. (a) Maximum-likelihood phylogeny of 50 copepod species inferred from 1,322 orthologous proteins. Nodes without labels have 100% bootstrap support. The scale bar indicates amino acid substitutions per site. (b–d) Maximum-likelihood ancestral trait reconstructions for mesopelagic habitat (b), diapause (c) and resting eggs (d), with pie charts showing the probability of trait presence at internal nodes.

Ancestral state reconstruction of traits indicates that calanoid ancestors colonised the mesopelagic zone (P=0.79) and acquired post-embryonic dormancy (P=0.97), whereas a shift towards embryonic dormancy occurred in Diaptomoidea ancestors associated with a transition to shallow-water habitats (P=0.97) and the emergence of resting eggs (P=0.98) (Fig. 1b,c,e Supplementary Table 4). The origin of DVM remains uncertain, but likely arose early in the evolutionary history of copepods (Fig. 1d). Our results suggest the following evolutionary scenario: mesopelagic colonisation and post-embryonic dormancy acquisition would have occurred in calanoid ancestors, followed by a shift towards embryonic dormancy and retreat from mesopelagic habitats in the Diaptomoidae. This scenario contrasts with the Bradford-Grieve hypothesis, that was based on morphological characters and fewer molecular markers phylogenies, and which proposed independent mesopelagic colonisations by epipelagic ancestors of Metridinidae and Pancalanina, with a habitat shift from benthic to epipelagic in Diaptomidae^20^.

However, it is worth noting that several taxa, particularly within the Paracalanidae and the Bradfordian families (that do not perform post-embryonic or embryonic dormancy) are not represented, nor are the hyperbenthic or anchialine cave taxa groups. Depending on the phylogenetic position of these groups within or outside the Pancalanina, the inferred probability of an ancestral origin of post-embryonic dormancy in calanoids could decrease, instead supporting multiple independent acquisitions. Therefore, the inference of an ancestral origin of post-embryonic dormancy should be considered provisional and contingent on current taxon sampling. Additional transcriptome sequencing of non-diapausing calanoid lineages will therefore be essential to clarify the evolutionary transitions of habitat and dormancy strategies within Calanoida.

### Calibrated time tree and oceanic palaeoclimate events

The calibrated time tree dated the origin of the calanoid stem lineage to 516.6 Ma (95% HPD = 349–666 Ma), whereas the most recent common ancestor of sampled calanoids was dated to 384 Ma (95% HPD = 234–495 Ma). Ancestral-state reconstruction indicated that mesopelagic colonisation and post-embryonic dormancy were already present in the most recent common ancestor of Calanoida and were therefore inferred to have been acquired along the calanoid stem lineage, between 516.6 and 384.6 Ma. This inferred interval encompasses the Ordovician episode of deep-ocean oxygenation (ca. 460–440 Ma)^22^ (Fig. 2a,b). We inferred a loss of post-embryonic dormancy in Diaptomoidea at 330 Ma (95% HPD = 200–429 Ma). Although the 95% HPD is broad, the posterior mean falls within the Carboniferous deep-sea deoxygenation interval (ca. 340-320 Ma)^22^ (Fig. 2a,b).

**Fig. 2.**
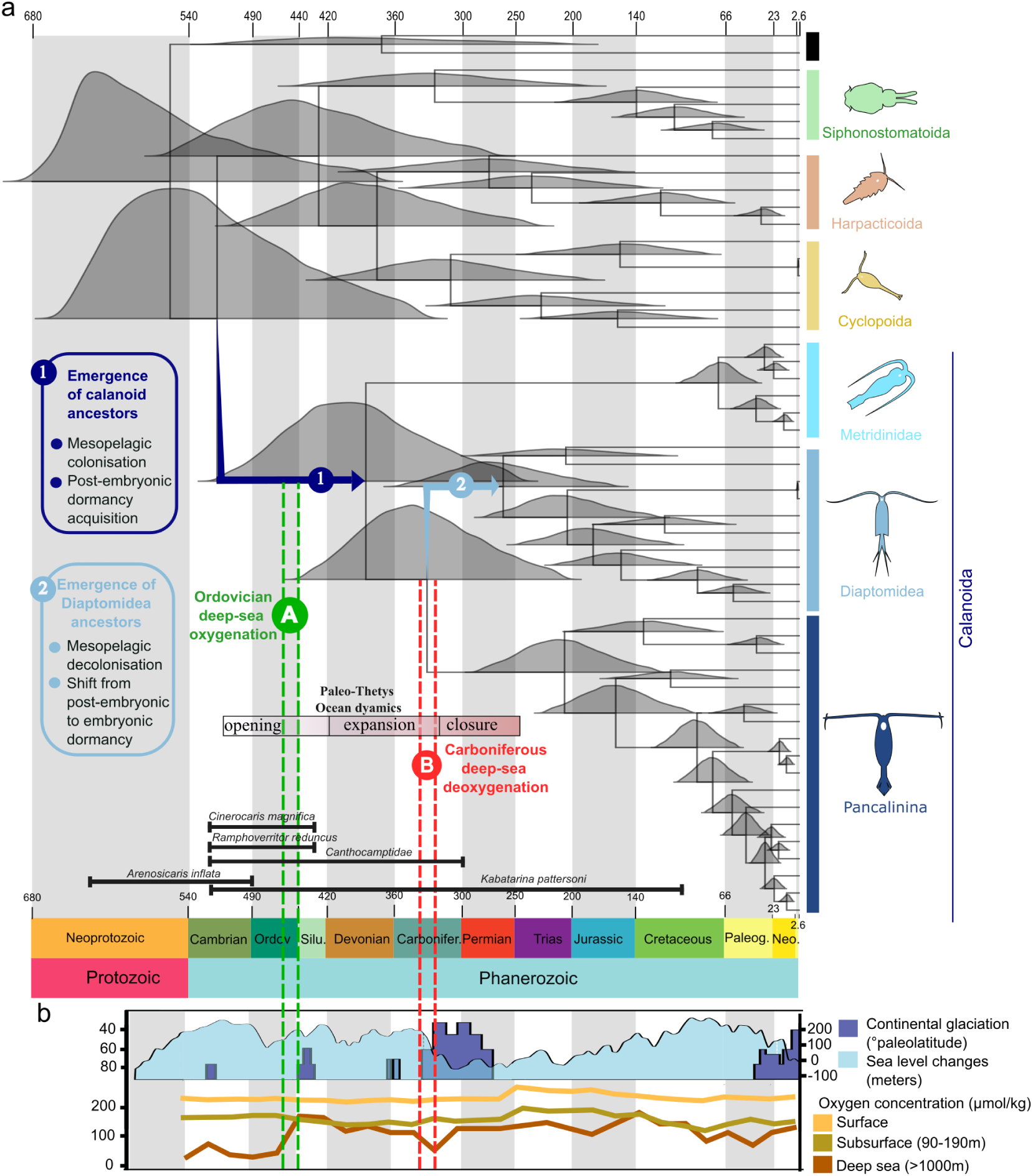
Copepod time tree and ocean palaeoclimate change. (a) Fossil-calibrated divergence time tree inferred with MCMCTree. The distribution in grey represents posterior estimates of divergence times across calibration replicates. The black bars represent the calibrated fossils. (b) Global palaeoclimate record of ocean oxygen dynamics (after Pohl et al., 2022^22^), highlighting two major deep-sea oxygenation/deoxygenation events noted A (green) and B (red).

The Ordovician deep-sea oxygenation was followed by the emergence and diversification of jawed vertebrates during the Silurian-Devonian period (440–360 Ma)^24^. Fish predation has been proposed to favour vertical displacement of copepods into deeper waters and the emergence of short-range vertical migration toward hypoxic layers^20^. Based on our time-calibrated phylogeny, calanoid ancestors are inferred to initiate differentiation in the late Cambrian. We propose that the combination of expanding habitable depth and increased predation pressure may have favoured the colonisation of mesopelagic habitats by calanoid ancestors through tracking of deeper oxyclines, potentially contributing to the emergence of long-range vertical migration. Increasing seasonality during the Late Devonian to Carboniferous (360–320 Ma)^25^, associated with glacial–interglacial cycles^26^, likely promoted lipid accumulation and extended hypoxia tolerance, setting the stage for the evolution of diapause as a long-term dormancy strategy in Pancalanina^27^.

The split between Metridinidae and the lineage leading to Diaptomoidea and Pancalanina was dated to 385 Ma (95% HPD = 234–495 Ma). The subsequent split between Diaptomoidea and Pancalanina was dated to 330 Ma (95% HPD = 200–429 Ma). The interval between these two splits occurred during the last Paleo-Tethys expansion in the Late Devonian-Carboniferous (440–320 Ma) and the establishment of mature ocean-basin structure^28^. During this period, the Paleo-Tethys Ocean underwent oceanographic differentiation, which was associated with major tectonic reorganisation, including the formation of deep basins and the establishment of distinct oceanic circulation systems^29,30^. Such features may act as physical barriers to dispersal by structuring water masses and limiting connectivity among planktonic populations^31^. As reduced connectivity is a key driver of geographic isolation and allopatric divergence^32^, we hypothesise that the expansion of the Paleo-Tethys Ocean may have contributed to lineage divergence by promoting spatial segregation of copepod populations. Under this scenario, Diaptomoidea and Pancalanina may have become partially isolated within the Paleo-Tethys basin, whereas basal lineages such as Arietelloidea (e.g., Metridinidae, which retain ancestral traits) persisted in the Panthalassa.

Geological evidence indicates that the Carboniferous–Permian (290–250 Ma) was characterised by pronounced sea-level fluctuations^33^ and widespread deoxygenation/reoxygenation in both deep and marginal marine environments^34^, particularly within the Paleo-Tethys Ocean^22^. In coastal areas, repeated transgressive–regressive cycles promote the formation of oxygen-depleted sediments^35^. In Diaptomidae, resting eggs sink and accumulate in sediments, where they remain viable under low-oxygen conditions^36^, and hypoxia can directly trigger development arrest of Diaptomidae eggs^37^. Although no fossil evidence currently constrains the past geographic distribution of these taxonomic groups, the timing of molecular phylogenetic divergence coincides with the Paleo-Tethys oceanic closure and deoxygenation. Therefore, we hypothesise that during the Carboniferous–Permian period, the combination of mesopelagic deoxygenation and sediment-associated hypoxia in neritic environments may have favoured a habitat shift in Diaptomoidae. Under these conditions, resting egg production may have been selectively favoured as a survival strategy, enabling persistence through episodic hypoxia and environmental instability and replacing post-embryonic dormancy.

### Expansion and collapse of gene families in marine copepods

As predicted proteins derived from transcriptomes may still contain assembly artefacts, we interpreted gene family evolution in terms of relative patterns of expansion and contraction across lineages and functional categories, rather than absolute gene copy numbers. The evolution of gene families (orthogroups, OGs) reveals a striking asymmetry in duplication and loss events. Siphonostomatoid copepods, which are predominantly parasitic, exhibit a pronounced contraction of gene families, consistent with patterns of genome reduction commonly observed in parasitic lineages^38–40^. In contrast, calanoid copepods display extensive lineage-specific expansions. A total of 900 OGs (722 genes) underwent expansion in calanoid ancestors, while 1,620 OGs experienced contraction in Diaptomoidae. Among these, 162 OGs (390 genes) correspond to secondary losses of gene families that had previously expanded in Calanoida ancestors (Fig. 3a). An alternative hypothesis to explain gene expansion in Calanoida ancestors would be whole-genome duplication (WGD). However, WGD has not been reported in copepods and the latest high quality genome sequence of *Eurytemora carolleeae* did not show any evidence of WGD^41^. Functional enrichment analysis of OGs showing the gain–loss dynamic identified significant over-representation of pathways involved in hypoxia response, including HIF-1 signaling, mitophagy, and MAPK (mitogen-activated protein kinase) signaling, as well as metabolic processes related to glutathione biosynthesis, lipid metabolism, and amino acid metabolism (Fisher exact test; FDR < 0.05; Fig. 3b).

**Fig. 3.**
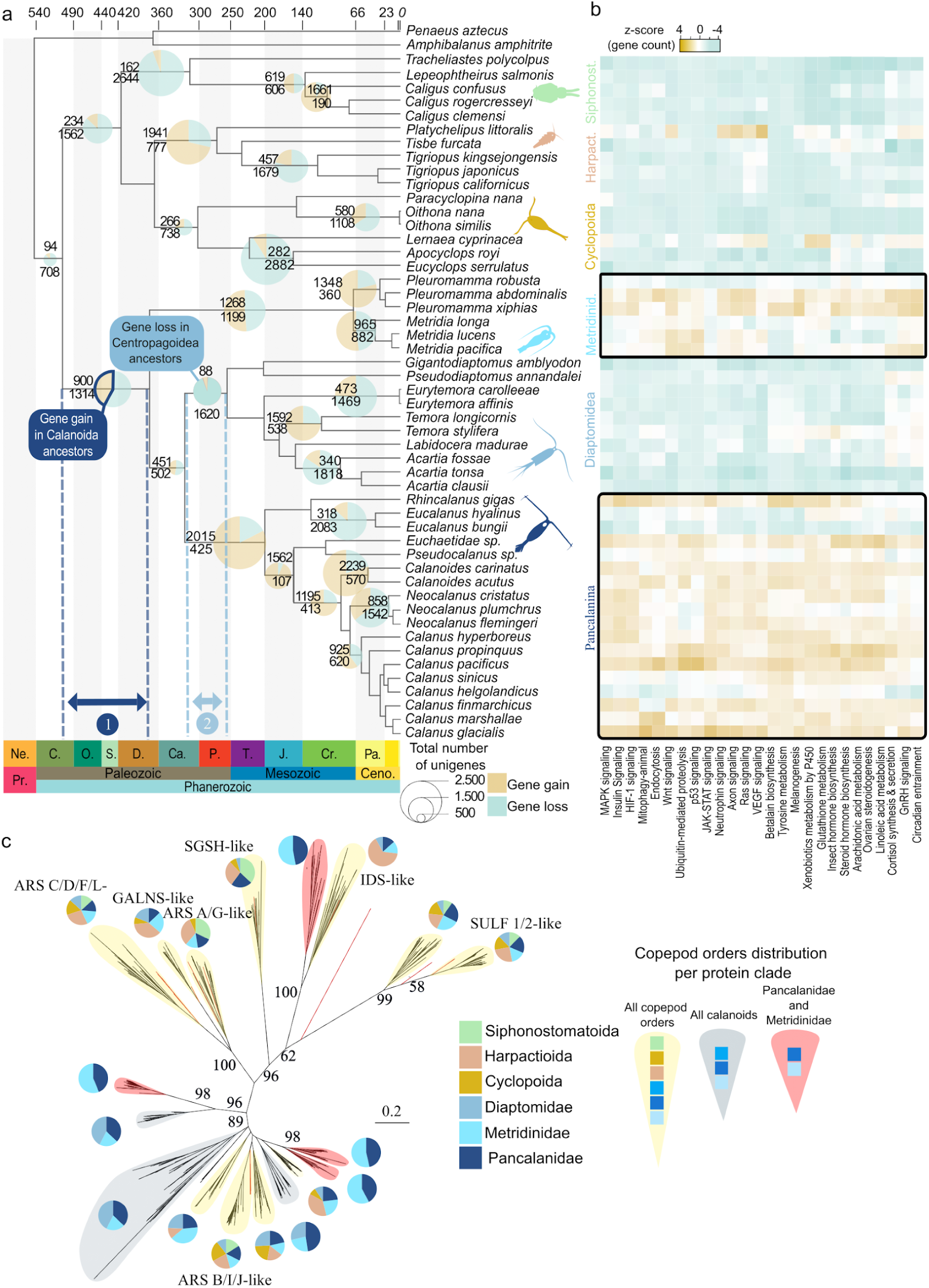
Gene family expansion and contraction in marine copepods. (a) Contraction and expansion of gene families inferred with CAFE. Pie charts at selected nodes indicate the proportion of gene loss (red) and gain (green), with size proportional to the number of affected genes. (b) Heatmap of gene count z-scores in KEGG pathways enriched among gene families showing the gain/loss pattern in Calanoida. (c) Maximum-likelihood phylogeny of arylsulfatase proteins. Pie charts indicate the distribution of proteins across copepod orders, with protein clades exhibiting gene-loss patterns highlighted in red.

The evolutionary pattern of gene expansion and loss within hypoxia-response pathways is exemplified by the phylogeny of arylsulfatases, enzymes involved in glycosaminoglycan sulfation and hypoxia response^42^ (Fig. 3c). While ten protein clades include representatives from all copepod orders, two protein clades are restricted to calanoids and four are specific to Pancalanina and Metridinidae, suggesting ancestral duplications in the calanoid lineage followed by secondary losses in Diaptomoidae.

The pattern of gene gains related to hypoxia in calanoid ancestors, followed by secondary losses in Diaptomidae, as illustrated by the phylogeny of arylsulfatases, is a more parsimonious interpretation of the observed duplication–loss dynamics (Fig. 3.b) than multiple independent acquisitions. This pattern also mirrors the inferred gain–loss of hypoxia-associated traits (Fig. 1b-e). We hypothesised that mesopelagic colonisation and exposure to hypoxia during dormancy and diel vertical migration were accompanied and may have been facilitated by the duplication of genes involved in hypoxia tolerance. We further propose that the loss of these genes in Diaptomidae is associated with a habitat shift toward oxygen-rich shallow waters, where selective pressures on hypoxia-related pathways were relaxed.

### Transcriptomic response to hypoxia in *Eucalanus hyalinus* from Oxygen Minimum Zone

The identification of gene family expansions in calanoid copepods revealed enrichment in pathways predicted to be associated with hypoxia response. While this pattern suggests that gene duplication may have contributed to hypoxia adaptation, it does not establish whether these genes are functionally involved in the physiological response to low oxygen. To bridge this gap between macroevolutionary patterns and present-day functions, we investigated the transcriptomic response of the hypoxia-tolerant copepod *Eucalanus hyalinus* in a natural OMZ. The Benguela upwelling off the Namibian coast provides a particularly relevant ecological setting, where strong vertical oxygen gradients and recurrent OMZ formation expose mesopelagic copepods to hypoxic conditions. *E. hyalinus* CV copepodites (stage V juveniles) were sampled *in situ* at the surface and within the OMZ. It allowed us to examine whether the gene families inferred to have expanded during calanoid evolution are associated with differential transcriptional responses under contemporary low-oxygen conditions.

Transcriptomic analysis revealed a significant up-regulation of 1,446 genes (436 with functional annotation) and down-regulation of 459 genes (43 with at least one functional annotation) in OMZ (|log_2_ fold change| ≥ 2 and FDR < 0.05; Fig. 4a). Differentially overexpressed genes in OMZ were tested for enrichment using KEGG pathway annotations, and significantly enriched pathways related to proteolysis, metabolite transport, amino acid metabolism, and glutathione metabolism (Hypergeometric test with FDR <0.05; Fig. 4b). Significantly overexpressed genes in OMZ that could be placed within metabolic or signaling pathways were further investigated.

**Fig. 4.**
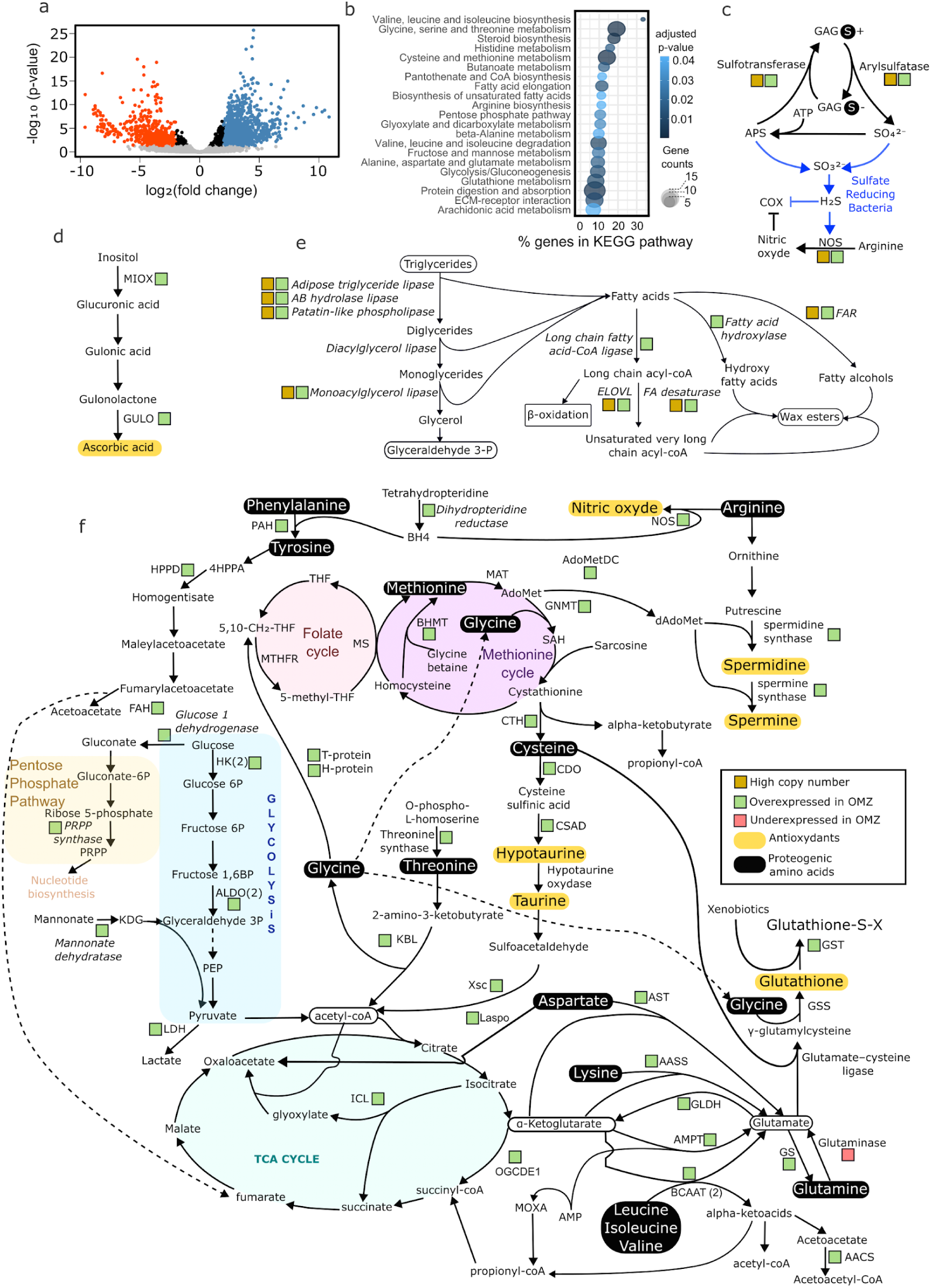
Transcriptomic response of *Eucalanus hyalinus* to oxygen minimum zone conditions compared with surface waters. (a) Volcano plot of differential gene expression. Genes with an adjusted p-value < 0.01 are considered significant; down-regulated genes (log_2_ fold change < –2) are shown in red, and up-regulated genes (log_2_ fold change > 2) in blue. (b) KEGG pathway enrichment analysis of differentially expressed genes. (c) Glycosaminoglycan metabolism pathway. (d) Ascorbic acid biosynthesis pathway. (e) Lipid biosynthesis pathway. (f) Amino acid metabolism pathway. Enzymes upregulated in hypoxia are represented with a green square.

*Eucalanus hyalinus* exhibits a coordinated ensemble of metabolic and cellular processes that collectively support survival in OMZ (Supplementary Results). These processes include developmental arrest, extracellular matrix (ECM) remodelling, sulfation and desulfation of ECM-associated glycosaminoglycans (Fig. 4.c), upregulation of metabolite transporters, lipid storage shift from triglycerides to wax esters (Fig. 4.e), rewiring of amino acid metabolism to sustain anaerobic ATP and antioxidant production (Fig. 4.d,f), and immune activation. Taken together, these results illustrate both evolutionary convergence between OMZ living copepods and hypoxia-adapted marine animals with the glycosaminoglycan (GAG) and sulphur metabolism^43,44^ and copepod-specific innovation through horizontal gene transfer with the glyoxylate shunt (Supplementary Results). These pathways are also present and activated under hypoxia in non-hypoxiphilic species, illustrating that it is the scale, efficiency and coordination of the response rather than the presence of individual pathways that constitutes a major determinant of OMZ adaptation in *E. hyalinus*^48^. Also, it is important to point out that surface waters and OMZ differ in several co-varying environmental parameters, including temperature (ΔT=6.5°C), hydrostatic pressure, and depth-associated microorganisms, which may also contribute to the observed transcriptional differences.

### Evolution of hypoxia-response genes in Calanoida

Orthogroups containing OMZ-upregulated genes in *E. hyalinus* were classified according to their biological processes based on integrated functional annotation, combining Pfam domain predictions, BLAST homology, KEGG pathway assignments, and manual curation (Fig. 5a,b). A significant increase in paralogues number in OGs associated to hypoxia response was observed in Metridinidae and Pancalanina compared with Diaptomoidae, particularly in families associated with oxidative stress, amino acid metabolism, and immune system (U-test, P<.05) (Fig. 5c). Paralogous expansions in these OGs harbours the same gain and loss pattern previously described at the genome scale (Fig. 2a). Among 15,402 OGs identified in *E. hyalinus*, 497 displayed the gain/loss profile and 368 were upregulated in OMZ. Fifty-four OGs belonged to both categories, significantly exceeding the 11.9 value expected under random association (odds ratio = 5.66, Fisher’s exact test, *P* = 4.8 × 10^−21^). This enrichment indicates that OMZ-induced genes are preferentially associated with orthogroups that expanded in calanoid ancestors and were secondarily lost in Diaptomidae.

**Fig. 5.**
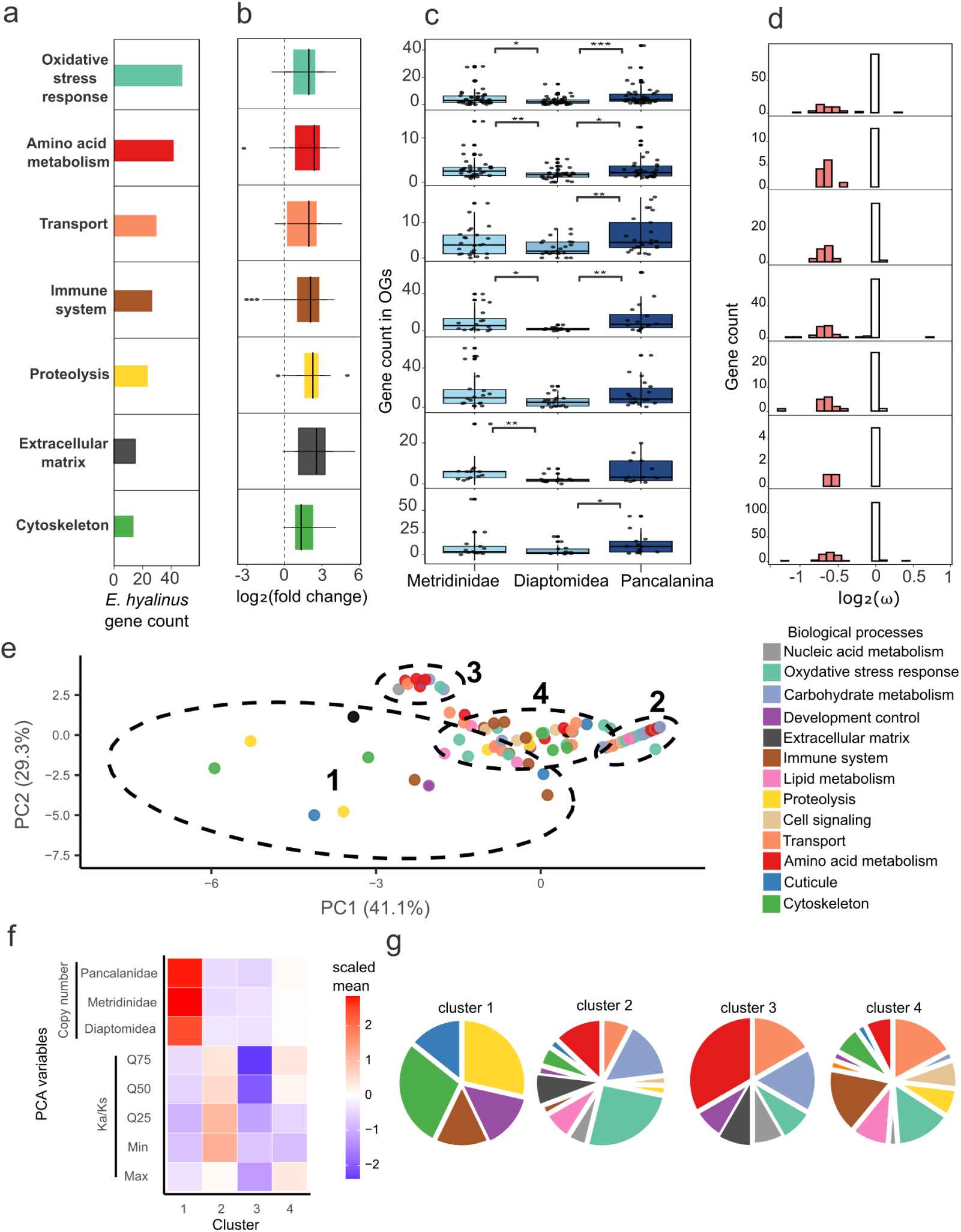
Evolutionary dynamics of hypoxia-induced genes related to their biological function. (a) Number of hypoxia-upregulated genes categorized by biological function in *E. hyalinus*. (b) Distribution of log_2_ fold change values. (c) Variation in gene family size across calanoid taxa by biological function (p < 0.05, **; p < 0.01, ***; p < 0.001). (d) Distribution of log_2_(*ω*) ratios by biological function. (e) PCA and k-mean clustering of orthogroups (OGs) by evolutionary variables (log_2_(*ω*) and lineage-specific copy number). (f) Contribution of evolutionary variables used in PCA to each k-mean cluster. (g) Distribution of the functional annotation of hypoxia induced genes in each cluster.

To identify patterns of selection and duplication associated with the functional roles of hypoxia-induced genes, we performed a principal component analysis (PCA) combining the distribution parameters of *ω*, the ratio between pairwise nonsynonymous (Ka) and synonymous (Ks) substitution rate and copy number variation of OGs (Fig. 5d,e). The first axis separates the OGs according to *ω* values and the second axis along copy number variations. K-means clustering (k=4) allows to differentiate OGs with high copy number and low *ω* in the Cluster 1, that contained only seven OGs related to proteolysis (serine protease and carboxypeptidase), immune system (Tumor Necrosis Factor), and cytoskeleton (myosin-motor domain-containing protein). Their *ω* profile (Fig. 5f) indicates that most genes evolve under strong purifying selection, supporting a dosage-driven mechanism^45^ that may enhance protein degradation, cytoskeleton maintenance and immune system. Cluster 2 grouped 38 OGs with moderate copy number and *ω*=1, pointing to neutral evolution and contained a large fraction of oxidative response genes. OGs with low copy number and *ω* were grouped into Cluster 3 that contained a large proportion of amino acid metabolism-related OGs (Fig. 5g). These families were typically small, often containing only a single gene upregulated under hypoxia and under purifying evolution. This pattern is consistent with subfunctionalisation, whereby one paralogue can be specialised for hypoxia-specific expression (Fig. 5f-g). Cluster 4 showed an intermediate profile, with moderate copy numbers and *ω* ≤1 and contains the largest amount of OGs, related to various biological processes.

### Conclusion

Our results indicate that hypoxia adaptation in calanoid copepods reflects the combined influence of palaeoclimate, trait emergence, and gene family expansion. Altogether, they allow a reinterpretation of the Bradford-Grieve hypothesis, which proposed that calanoid copepods colonised the pelagic realm through successive invasions linked to major paleoceanographic transitions, with early lineages entering the water column during the Devonian and later radiations associated with deep-ocean ventilation and subsequent deoxygenation events. According to our new results, we hypothesise that during the Ordovician deep-sea oxygenation, intensified visual predation by vertebrates in surface layers likely forced calanoid ancestors to track the deepening oxycline, promoting colonisation of the mesopelagic zone, the acquisition of post-embryonic dormancy, and the expansion of hypoxia-response gene families. Later climatic fluctuations, notably the Carboniferous deoxygenation–reoxygenation, drove divergent trajectories: Metridinidae and Pancalanina retained post-embryonic dormancy and paralogue expansions, whereas Diaptomoidea probably retreated from mesopelagic to shallow habitats, shifted from post-embryonic to embryonic dormancy, and exhibited extensive gene family contractions. Nevertheless, the current taxon sampling remains biased toward diapausing calanoid lineages. Additional transcriptome sequencing of non-diapausing calanoid taxa, particularly within the Paracalanidae and Bradfordian families, will be essential to confirm the evolutionary transitions in habitat and dormancy strategies inferred here. The transcriptomic response of *E. hyalinus* in the Benguela OMZ illustrates how these paralogues are mobilised under contemporary hypoxia, coupling developmental arrest, extracellular matrix catabolism, sulphur metabolism, antioxidant production, and lipid-to-wax ester conversion. Evolutionary analyses further reveal that proteolysis and transport protein families expanded under purifying selection, suggesting dosage effects.

## Materials and Methods

### Copepod molecular database

A total of 50 copepod transcriptomes were analysed in this study (Supplementary Table 1), including six newly assembled datasets and 44 reassembled from previously published sources. Two outgroup taxa were included for phylogenetic analyses: the malacostracan *Penaeus aztecus* and the thecostracan *Amphibalanus amphitrite*. Outgroups were selected based on the availability of high-quality transcriptomic resources and fossil calibrations suitable for divergence-time estimation. These two taxa are relatively distant from Copepoda, and the use of closer outgroups (e.g., Branchiopoda) may improve phylogenetic resolution in future analyses^46,47^.

### Sub-arctic sampling

Adult females of *Calanus hyperboreus*, *Pseudocalanus* sp., and *Metridia longa* were collected in December 2019 at the Rimouski monitoring station of the Atlantic Zone Monitoring Program (AZMP), located in the St. Lawrence maritime estuary (48°40′ N, 68°35′ W) at a depth of 320 m. Total RNA was extracted using the NucleoSpin RNA XS kit (Macherey-Nagel), and cDNA was synthesized with the Illumina SMARTer Ultra Low RNA kit. Paired-end libraries were prepared using the NEBNext DNA Sample Prep Reagent Set (Ozyme) and sequenced on an Illumina MiSeq platform.

### OMZ sampling

*Eucalanus hyalinus, Calanoides carinatus and Pleuromamma robusta* sampling was conducted off the Namibian coast during the D356 cruise of the RRS Discovery. In October 2010, a surface sample was collected using a WP2-net drifting at a depth of 10 meters for 20 minutes at coordinates 23.001°S, 13.055°E (Supplementary Table 7). The average temperature, salinity, and oxygen concentration at this site were 16.4°C, 35.6 g/kg, and 226.9 µmol/kg, respectively. In October 2010, an OMZ sample was taken from a depth of 250 to 400 meters at coordinates 18.993°S, 10.335°E (Supplementary Table 7). The average temperature, salinity, and oxygen concentration at this site were 9.9°C, 34.9 g/kg, and 44.8 µmol/kg, respectively. All samples were immediately fixed with a saturated ammonium sulphate solution either, for surface samples upon recovery or for mesopelagic samples using a custom-built in-situ fixation unit. Samples were then stored at 8°C for 24 hours and afterward stages CIII to CV were isolated and stored at -20°C in a fresh ammonium sulphate solution. Total RNA was isolated from CV stage specimens using the trizol reagent, with four replicates for each condition. Following RNA quality control, paired-end sequencing (100 bp) was performed using HiSeq2000. For *E. hyalinus* six samples were kept for differential expression analysis: three from the oxygenated surface zone and three from the OMZ. Only one sample for *C. carinatus* and one for *P. robusta* were kept.

### Transcriptome assembly, proteome prediction and annotation

Raw sequencing files were retrieved from the NCBI database using the SRA Toolkit v2.11 v2.11 (https://github.com/ncbi/sra-tools). Read quality was assessed with FastQC v0.11.9^48^ and adapter and low-quality regions were trimmed using BBmap tool v38.90 (https://sourceforge.net/projects/bbmap/). New and downloaded transcriptomic data were assembled with Trinity v2.8.4 using default parameters^49^. Assembly completeness was evaluated with BUSCO v5.2.2 against the Arthropoda ortholog database (odb10, 09/2020)^50^ and assemblies with ≤25% missing BUSCOs were retained for downstream analyses. Protein-coding sequences were predicted with TransDecoder v5.5.0 and clustered using CD-HIT v4.8.1^51^ (parameters: *-c 0.95 -G 0 -aS 0.7*). The same workflow was applied to the six newly assembled transcriptomes. To evaluate whether variation in predicted protein counts was driven by assembly completeness, we fitted ordinary least squares linear models testing the effect of BUSCO completeness on predicted protein number. The significance of predictors was assessed using two-sided t-tests on regression coefficients. BUSCO completeness was not significantly associated with predicted protein counts (R²=0.15, p=0.24)

Functional annotation of all predicted copepod proteomes followed a multistep pipeline. InterProScan v.5.54-87^52^ was used to detect conserved domains and assign Pfam, Gene Ontology, and InterPro annotations^53^. EggNOG-mapper v2.1.9 was then used to assign orthology and KEGG pathway information based on the eggNOG v5.0.2 database^54,55^. Finally, DIAMOND v2.0.4 was employed to perform BLASTP searches against the UniProtKB and *Drosophila melanogaster* UniProt reference proteomes^56^.

### Phylogenomic analyses

Putative orthologs were identified using the Reciprocal Best Hits (RBH) approach implemented with DIAMOND v2.0.2, run in *very sensitive* mode and limited to one best target sequence per query^57^. After reciprocal pairs were identified, three quality filters were applied to retain high-confidence orthologs: a minimum of 30% sequence identity, more than 60% coverage of the shorter sequence, and an e-value below 10^−6^. Only RBHs shared by all proteomes were retained for downstream analyses. Protein sequences were aligned with MAFFT v7.475 using automatic parameter selection, and poorly aligned regions were removed with trimAl v1.2, applying a gap threshold of 0.9, a similarity threshold of 0.001, a window size of three residues, and a minimum consensus requirement of 25% ^58,59^. Each protein was assigned its own best-fit substitution model determined by ModelTest-NG^60^. The resulting alignments were concatenated and used to infer a partitioned maximum likelihood phylogeny in RAxML-NG v1.1.0 with 500 standard bootstrap replicates^61^. This phylogenomic pipeline was applied to the dataset of 50 copepod species and two outgroup taxa and yielded 1,322 orthologous groups. The final concatenated alignment comprised 494,348 amino acid positions with 294,154 (59.5%) parsimony-informative sites and 5.6% of gaps. The phylogenetic tree was visualised using FigTree v1.4.4 (http://tree.bio.ed.ac.uk/software/figtree) and graphically refined with Inkscape v1.3.2 (https://inkscape.org).

### Ancestral trait reconstruction

A table summarizing four ecological and physiological traits (mesopelagic colonisation, diapause, resting eggs and DVM) was compiled from the literature (Supplementary Table 2) and a binary presence–absence matrix was generated. For DVM, only the presence or absence of diel vertical migration was considered; no distinction was made between weak and strong migratory amplitudes. We defined ‘diapause’ in a broad physiological sense as a reversible period of metabolic suppression and reduced activity, typically associated with lipid accumulation and ontogenetic vertical descent to deep waters. This definition encompasses both the facultative quiescence observed in *Metridinidae* and the hormonally programmed developmental arrest of *Calanidae* and *Eucalanidae*. Because trait definitions and ecological observations are not always consistent across the literature, species showing contradictory or uncertain trait descriptions were annotated and scored as undefined in the matrix to avoid introducing bias in the ancestral trait reconstruction. The evolutionary history of each trait was reconstructed from this matrix and the phylogenetic tree using maximum likelihood (ML) and asymmetrical Mk models implemented in Mesquite v3.81^62^. Bayesian reconstruction was performed using the MultiState model implemented in BayesTraits v5.0.1^63^.

### Estimation of species divergence time

Divergence times were estimated from the concatenated alignment of 1,322 single-copy orthologous proteins using the Bayesian relaxed molecular clock implemented in MCMCTree from the PAML package v4.9^64^. The maximum likelihood topology previously inferred was used as the fixed input tree. Five fossil calibration points were applied according to Wolfe *et al.*^65^ (Supplementary Table 8). Model selection was performed in MEGA11^66^ based on the Bayesian Information Criterion (BIC) and Akaike Information Criterion (AIC), identifying the LG substitution model as optimal. The Hessian matrix was first computed under this model to estimate the gradient and variance–covariance parameters. The Markov chain Monte Carlo (MCMC) analysis was run for 200 million generations, with the first 50% discarded as burn-in, and posterior samples collected every 500 generations. Three independent runs were performed from different starting seeds. Convergence among the three independent runs was evaluated using the Gelman-Rubin potential scale reduction factor (PSRF) and by the effective sample size (ESS). All PSFR values were below 1.05 and all ESS below 200, indicating good mixing and convergence among independent runs. Convergence was additionally evaluated by comparing posterior mean node ages and their 95% HPD intervals across runs (Supplementary Fig. 3) and were highly consistent across runs, supporting the robustness and reliability of the divergence time estimates. Resulting chronograms were visualised using the R package *MCMCtreeR* v1.1^67^. Effective prior and posterior node-age distributions were compared using MCMCTree runs performed without and with sequence data. Posterior 95% HPD intervals were consistently narrower than effective prior HPDs across all nodes (mean posterior/prior HPD width ratio = 0.270; median = 0.256; range = 0.003–0.799), corresponding to an average 73.0% reduction in uncertainty (Supplementary Fig. 4).

### Gain and Loss of orthologous genes

Gene family inference was performed using OrthoFinder v2.5.2^68^ on the predicted proteomes of 53 species. The analysis identified 143,524 orthogroups (OGs), each representing a set of orthologous and paralogous genes derived from a common ancestral locus. Two filtering criteria were applied prior to downstream analyses: (i) removal of OGs containing >100 protein sequences in any single species, and (ii) exclusion of species-specific OGs present in only one taxon. The evolution of gene family size was then analysed using CAFE v5^69^ with parameters “-k 2 -p,” based on the filtered OG dataset and an ultrametric species tree. The global distribution of gene family gains and losses across the phylogeny was summarized with CafePlotter v0.2.0 (https://github.com/moshi4/CafePlotter). To identify lineage-specific gene expansions, for each orthogroup, two preliminary tests were used: the Shapiro–Wilk test to assess normality of gene count distributions and the Bartlett test to evaluate variance homogeneity among groups. OGs failing either test (p < 0.01 for Shapiro–Wilk; p < 0.05 for Bartlett) were excluded. The remaining OGs were analysed using a Welch’s t-test comparing mean gene counts in Metridinidae or Pancalanina against other copepod orders and superfamilies. In total, 381 orthogroups (<0.3%) displayed a significant difference (p < 0.05), indicating lineage-specific gene expansion or duplication. To examine the functional composition of expanded gene families, z-scores of gene occurrence were calculated across species for KEGG pathways and visualised with heatmaps^70–72^.

#### Gene phylogeny

For the phylogenetic analysis of the arylsulfatase, the protein sequences were identified based on the presence of the complete arylsulfatase domain. A total of 519 proteins were detected across all copepod-predicted proteomes (length >300 amino acids; e-value <10^−6^). After adding fourteen human arylsulfatase reference sequences, a 245-amino-acid alignment was generated using MUSCLE^73^, and a maximum-likelihood phylogenetic tree was constructed in MEGA11^66^ under the WAG+F substitution model with 100 bootstrap replicates. For the isocitrate lyase, sequences identified in copepod proteomes were used as a query against the NCBI nr database to retrieve all the closest homologs. All sequences were aligned with MUSCLE (default parameters), and conserved blocks were manually curated to remove ambiguously aligned regions. The final alignment was used to infer a maximum likelihood tree in MEGA11 under the LG + I + G substitution model with 100 bootstrap replicates.

### Differential expression analysis in Eucalanus hyalinus between surface and oxygen minimum zone

RNA-seq reads were mapped against the *E. hyalinus* transcriptome using bwa mem^74^ and duplicate reads were removed with samtools^75^. Read counts were extracted and normalized using EdgeR v4.0.16^72,76^. MA-plot, displaying a log ratio (M) vs. an average (A) in order to visualize the differences between two groups, confirmed globally homogeneous gene expression between normoxic and hypoxic conditions (Supplementary Fig. 2). Differential expression analysis was performed between normoxia and hypoxia treatments, and genes with a Benjamini–Hochberg adjusted *p*-value ≤ 0.05 were considered significantly differentially expressed. Functional enrichment of differentially expressed genes was assessed using KEGG pathway analysis based on Fisher’s exact test, with significant pathways retained at FDR < 0.05. A total of 436 up-regulated genes were manually annotated by protein domain inspection and BLAST searches against the NCBI nr database, and subsequently classified into 13 functional categories (Supplementary Table 5).

### Evolutionary patterns of hypoxia-response genes

Significance of differences in orthogroup size distributions among copepod lineages was assessed using a Wilcoxon rank-sum test, allowing the comparison of gene family expansion levels between Metridinidae, Pancalanina, and Diaptomoidae. To characterize selective pressures acting after gene duplication, pairwise nonsynonymous (Ka) and synonymous (Ks) substitution rates were estimated between paralogous genes within each orthogroup (OG) of *E. hyalinus* using SeqinR v4.2^71,72,76,77^. For each OG containing at least two paralogues, amino acid sequences were aligned and the corresponding coding DNA sequences were back-translated to generate codon-preserving alignments. Pairwise Ka and Ks values were then computed using a codon-based substitution model, and the resulting Ka/Ks ratios (*ω*) were assembled into matrices from which undefined or infinite values were removed. To explore relationships between selective regimes and gene family expansion, a principal component analysis (PCA) was performed on OG-level summary statistics (*ω* quantiles and copy number in the three taxonomic groups) after z-score normalization. Orthogroups were partitioned into four clusters using k-means clustering (k = 4) based on the first three PCA axes that contained more than 85% of the variance. To assess the relative contribution of each variable across clusters, *ω* summary statistics and copy numbers were standardized and averaged within clusters and visualized as a heatmap. All data and codes used in this study are available at https://github.com/madoui/Hypoxia.

## Acknowledgements

We thank the Agence Nationale de la Recherche for funding this study under grant ANR-22-CE02-0023 (TRAITZOO project). This work also benefited from the support of the Institut des Sciences du Calcul et des Données (ISCD) at Sorbonne Université (SU), through the junior research team FORMAL (From Observing to Modeling Ocean Life). RK acknowledges support via the DFG funded Cluster of Excellence “Future Ocean”; project CP0923. We gratefully thank the Genouest bioinformatic platform and Olivier Colin from Rennes, France and Genoscope from Evry, France for IT and sequencing support. We are grateful to Stéphane Plourde and Geneviève Perrin at Maurice-Lamontagne Institute, Fisheries and Oceans Canada, for graciously providing the St. Lawrence estuary samples. We acknowledge the RRS Discovery crew members. Co-authors wish to thank French and European public taxpayers who fund their salaries.

## References

1. Wright, J. J., Konwar, K. M. & Hallam, S. J. Microbial ecology of expanding oxygen minimum zones. Nat Rev Microbiol 10, 381–394 (2012).

2. Ulloa, O., Canfield, D. E., DeLong, E. F., Letelier, R. M. & Stewart, F. J. Microbial oceanography of anoxic oxygen minimum zones. Proc. Natl. Acad. Sci. U.S.A. 109, 15996–16003 (2012).

3. Breitburg, D. et al. Declining oxygen in the global ocean and coastal waters. Science 359, eaam7240 (2018).

4. Stramma, L., Johnson, G. C., Sprintall, J. & Mohrholz, V. Expanding Oxygen-Minimum Zones in the Tropical Oceans. Science 320, 655–658 (2008).

5. Steinberg, D. K. & Landry, M. R. Zooplankton and the Ocean Carbon Cycle. Annual Review of Marine Science 9, 413–444 (2017).

6. Dam, H. G. Evolutionary Adaptation of Marine Zooplankton to Global Change. Annual Review of Marine Science 5, 349–370 (2013).

7. Wishner, K. F., Seibel, B. & Outram, D. Ocean deoxygenation and copepods: coping with oxygen minimum zone variability. Biogeosciences 17, 2315–2339 (2020).

8. Dahms, H.-U. Dormancy in the Copepoda — an overview. Hydrobiologia 306, 199–211 (1995).

9. Baumgartner, M. F. & Tarrant, A. M. The Physiology and Ecology of Diapause in Marine Copepods. Annu. Rev. Mar. Sci. 9, 387–411 (2017).

10. Bandara, K., Varpe, Ø., Wijewardene, L., Tverberg, V. & Eiane, K. Two hundred years of zooplankton vertical migration research. Biological Reviews 96, 1547–1589 (2021).

11. Roman, M. R., Brandt, S. B., Houde, E. D. & Pierson, J. J. Interactive Effects of Hypoxia and Temperature on Coastal Pelagic Zooplankton and Fish. Front. Mar. Sci. 6, (2019).

12. Hochachka, P. W. & Lutz, P. L. Mechanism, origin, and evolution of anoxia tolerance in animals☆. Comparative Biochemistry and Physiology Part B: Biochemistry and Molecular Biology 130, 435–459 (2001).

13. Lee, P., Chandel, N. S. & Simon, M. C. Cellular adaptation to hypoxia through hypoxia inducible factors and beyond. Nat Rev Mol Cell Biol 21, 268–283 (2020).

14. Mills, D. B. et al. The last common ancestor of animals lacked the HIF pathway and respired in low-oxygen environments. eLife 7, e31176.

15. Graham, A. M. & Barreto, F. S. Loss of the HIF pathway in a widely distributed intertidal crustacean, the copepod Tigriopus californicus. Proceedings of the National Academy of Sciences 116, 12913–12918 (2019).

16. Graham, A. M. & Barreto, F. S. Independent Losses of the Hypoxia-Inducible Factor (HIF) Pathway within Crustacea. Molecular Biology and Evolution 37, 1342–1349 (2020).

17. Rytkönen, K. T. et al. Subfunctionalization of Cyprinid Hypoxia-Inducible Factors for Roles in Development and Oxygen Sensing. Evolution 67, 873–882 (2013).

18. Meng, J., Wang, T., Li, B., Li, L. & Zhang, G. Oxygen sensing and transcriptional regulation under hypoxia exposure in the mollusk *Crassostrea gigas*. Science of The Total Environment 853, 158557 (2022).

19. Yu, J. J. et al. Time Domains of Hypoxia Responses and -Omics Insights. Front. Physiol. 13, 885295 (2022).

20. Bradford-Grieve, J. M. Colonization of the pelagic realm by calanoid copepods. Hydrobiologia 485, 223–244 (2002).

21. Blanco-Bercial, L., Bradford-Grieve, J. & Bucklin, A. Molecular phylogeny of the Calanoida (Crustacea: Copepoda). Molecular Phylogenetics and Evolution 59, 103–113 (2011).

22. Pohl, A. et al. Continental configuration controls ocean oxygenation during the Phanerozoic. Nature 608, 523–527 (2022).

23. Berger, C. A., Steinberg, D. K., Copeman, L. A. & Tarrant, A. M. Comparative analysis of the molecular starvation response of Southern Ocean copepods. Molecular Ecology 34, e17371 (2025).

24. Brazeau, M. D. & Friedman, M. The origin and early phylogenetic history of jawed vertebrates. Nature 520, 490–497 (2015).

25. De Vleeschouwer, D. et al. Timing and pacing of the Late Devonian mass extinction event regulated by eccentricity and obliquity. Nat Commun 8, 2268 (2017).

26. Montañez, I. P. et al. Climate, pCO2 and terrestrial carbon cycle linkages during late Palaeozoic glacial–interglacial cycles. Nature Geosci 9, 824–828 (2016).

27. Cavallo, A. & Peck, L. S. Lipid storage patterns in marine copepods: environmental, ecological, and intrinsic drivers. ICES J Mar Sci 77, 1589–1601 (2020).

28. Stampfli, G. M. & Borel, G. D. A plate tectonic model for the Paleozoic and Mesozoic constrained by dynamic plate boundaries and restored synthetic oceanic isochrons. Earth and Planetary Science Letters 196, 17–33 (2002).

29. Berra, F. & Angiolini, L. The Evolution of the Tethys Region throughout the Phanerozoic: A Brief Tectonic Reconstruction. in 1–27 (2014). doi:10.1306/13431840M1063606.

30. Gérard, J. et al. Exploring the mechanisms of Devonian oceanic anoxia: impact of ocean dynamics, palaeogeography, and orbital forcing. Climate of the Past 21, 239–260 (2025).

31. Cowen, R. K. & Sponaugle, S. Larval dispersal and marine population connectivity. Ann Rev Mar Sci 1, 443–466 (2009).

32. Palumbi, S. R. Genetic Divergence, Reproductive Isolation, and Marine Speciation. Annual Review of Ecology and Systematics 25, 547–572 (1994).

33. Fielding, C. R., Frank, T. D. & Isbell, J. L. The late Paleozoic ice age—A review of current understanding and synthesis of global climate patterns. in Resolving the Late Paleozoic Ice Age in Time and Space (eds Fielding, C. R., Frank, T. D. & Isbell, J. L.) 0 (Geological Society of America, 2008). doi:10.1130/2008.2441(24).

34. Isozaki, Y. Permo-Triassic Boundary Superanoxia and Stratified Superocean: Records from Lost Deep Sea. Science 276, 235–238 (1997).

35. Tyson, R. V. & Pearson, T. H. Modern and ancient continental shelf anoxia: an overview. *Geological Society, London*, Special Publications 58, 1–24 (1991).

36. Marcus, N. H. Ecological and evolutionary significance of resting eggs in marine copepods: past, present, and future studies. Hydrobiologia 320, 141–152 (1996).

37. Jørgensen, T. S., Jepsen, P. M., Petersen, H. C. B., Friis, D. S. & Hansen, B. W. Eggs of the copepod Acartia tonsa Dana require hypoxic conditions to tolerate prolonged embryonic development arrest. BMC Ecol 19, 1 (2019).

38. Cavalier-Smith, T. Economy, Speed and Size Matter: Evolutionary Forces Driving Nuclear Genome Miniaturization and Expansion. Annals of Botany 95, 147–175 (2005).

39. Poulin, R. & Randhawa, H. S. Evolution of parasitism along convergent lines: from ecology to genomics. Parasitology 142, S6–S15 (2015).

40. Wolf, Y. I. & Koonin, E. V. Genome reduction as the dominant mode of evolution. BioEssays 35, 829–837 (2013).

41. Du, Z., Gelembiuk, G., Moss, W., Tritt, A. & Lee, C. E. The Genome Architecture of the Copepod *Eurytemora carolleeae —* the Highly Invasive Atlantic Clade of the *Eurytemora affinis* Species Complex. *Genomics*, Proteomics & Bioinformatics 22, qzae066 (2024).

42. Bhattacharyya, S. & Tobacman, J. K. Hypoxia Reduces Arylsulfatase B Activity and Silencing Arylsulfatase B Replicates and Mediates the Effects of Hypoxia. PLOS ONE 7, e33250 (2012).

43. Zhu, F.-C. et al. Insights into the strategy of micro-environmental adaptation: Transcriptomic analysis of two alvinocaridid shrimps at a hydrothermal vent. PLoS ONE 15, e0227587 (2020).

44. Gan, Z. et al. Comparative transcriptomic analysis of deep- and shallow-water barnacle species (Cirripedia, Poecilasmatidae) provides insights into deep-sea adaptation of sessile crustaceans. BMC Genomics 21, 240 (2020).

45. Kondrashov, F. A., Rogozin, I. B., Wolf, Y. I. & Koonin, E. V. Selection in the evolution of gene duplications. Genome Biol 3, research0008.1 (2002).

46. Bernot, J. P. et al. Major Revisions in Pancrustacean Phylogeny and Evidence of Sensitivity to Taxon Sampling. Mol Biol Evol 40, msad175 (2023).

47. Yu, H. Y., Chu, K. H., Tsang, L. M. & Ma, K. Y. Incomplete lineage sorting and long-branch attraction confound phylogenomic inference of Pancrustacea. Front. Ecol. Evol. 12, (2024).

48. Andrews, S. FastQC A Quality Control tool for High Throughput Sequence Data. (2010).

49. Grabherr, M. G. et al. Trinity: reconstructing a full-length transcriptome without a genome from {RNA}-Seq data. 29, 644.

50. Manni, M., Berkeley, M. R., Seppey, M., Simão, F. A. & Zdobnov, E. M. BUSCO Update: Novel and Streamlined Workflows along with Broader and Deeper Phylogenetic Coverage for Scoring of Eukaryotic, Prokaryotic, and Viral Genomes. Molecular Biology and Evolution 38, 4647–4654 (2021).

51. Fu, L., Niu, B., Zhu, Z., Wu, S. & Li, W. CD-HIT: accelerated for clustering the next-generation sequencing data. Bioinformatics 28, 3150–3152 (2012).

52. Jones, P. et al. InterProScan 5: genome-scale protein function classification. Bioinformatics (Oxford, England) 30, 1236–1240 (2014).

53. Mitchell, A., et al. The {InterPro} protein families database: the classification resource after 15 years. 43, D213–221.

54. Cantalapiedra, C. P., Hernández-Plaza, A., Letunic, I., Bork, P. & Huerta-Cepas, J. eggNOG-mapper v2: Functional Annotation, Orthology Assignments, and Domain Prediction at the Metagenomic Scale. Mol Biol Evol 38, 5825–5829 (2021).

55. Kanehisa, M., Goto, S., Sato, Y., Furumichi, M. & Tanabe, M. KEGG for integration and interpretation of large-scale molecular data sets. Nucleic acids research 40, D109–14 (2012).

56. Buchfink, B., Xie, C. & Huson, D. H. Fast and sensitive protein alignment using DIAMOND. Nature Methods 12, 59–60 (2015).

57. Buchfink, B., Reuter, K. & Drost, H.-G. Sensitive protein alignments at tree-of-life scale using DIAMOND. Nat Methods 18, 366–368 (2021).

58. Katoh, K. & Standley, D. M. MAFFT multiple sequence alignment software version 7: improvements in performance and usability. Molecular biology and evolution 30, 772–780 (2013).

59. Capella-Gutiérrez, S., Silla-Martínez, J. M. & Gabaldón, T. trimAl: a tool for automated alignment trimming in large-scale phylogenetic analyses. Bioinformatics 25, 1972–1973 (2009).

60. Darriba, D. et al. ModelTest-NG: A New and Scalable Tool for the Selection of DNA and Protein Evolutionary Models. Mol Biol Evol 37, 291–294 (2020).

61. Kozlov, A. M., Darriba, D., Flouri, T., Morel, B. & Stamatakis, A. RAxML-NG: a fast, scalable and user-friendly tool for maximum likelihood phylogenetic inference. Bioinformatics 35, 4453–4455 (2019).

62. Maddison, W. P. & Maddison, D. R. Mesquite: a modular system for evolutionary analysis.

63. Meade, A. & Pagel, M. Ancestral State Reconstruction Using BayesTraits. in Environmental Microbial Evolution: Methods and Protocols (ed. Luo, H.) 255–266 (Springer US, New York, NY, 2022). doi:10.1007/978-1-0716-2691-7_12.

64. Yang, Z. PAML 4: phylogenetic analysis by maximum likelihood. Mol Biol Evol 24, 1586–1591 (2007).

65. Wolfe, J. M., Daley, A. C., Legg, D. A. & Edgecombe, G. D. Fossil calibrations for the arthropod Tree of Life. Earth-Science Reviews 160, 43–110 (2016).

66. Tamura, K., Stecher, G. & Kumar, S. MEGA11: Molecular Evolutionary Genetics Analysis Version 11. Molecular Biology and Evolution 38, 3022–3027 (2021).

67. Puttick, M. N. MCMCtreeR: functions to prepare MCMCtree analyses and visualize posterior ages on trees. Bioinformatics 35, 5321–5322 (2019).

68. Emms, D. M. & Kelly, S. OrthoFinder: phylogenetic orthology inference for comparative genomics. Genome Biology 20, 238 (2019).

