## Supplementary files for "Hypoxia adaptation shapes genomic architecture and vertical niche transitions in copepods"

* corresponding author

**Competing Interest Statement:** Disclose any competing interests here.

**Supplementary Table 1: Transcriptomic read sets.** **2**
**Supplementary Table 2. Table of three key functional traits across copepod species and related studies.** **5**
**Supplementary Table 3. Table of references of the three key functional traits** **6**
**Supplementary Table 4: Ancestral trait reconstruction** **9**
**Supplementary Table 5. Classification of upregulated genes** **9**
**Supplementary Table 6. Enzyme symbols** **10**
**Supplementary Table 7: Copepod sampling for newly assembled transcriptomes.** **11**
**Supplementary Table 8 : Five fossil calibrations for arthropods from Wolfe et al. 2016** **11**
**Supplementary Figure 1: Maximum-likelihood phylogenetic tree of isocitrate lyase.** **12**
**Supplementary Figure 2: MA plot of Eucalanus hyalinus transcripts** **13**
**Supplementary Figure 3: Convergence analysis between the three independent MCMC analyses** **14**
**Supplementary Figure 4: MCMCTree prior vs posterior highest posterior density widths** **15**
**Supplementary Results: Transcriptomic response of Eucalanus hyalinus in OMZ** **16**
**References** **20**

#### Supplementary Table 1: Transcriptomic read sets.

| Order | Family | Organism | # predicted proteins after CDhits | BUSCO completeness + partial(%) | Source |
|---|---|---|---|---|---|
| Calanoida | **Acartiidae** | *Acartia clausi* | 23 426 | 94.5 | SRR20015063 |
| | | *Acartia fossae* | 30 212 | 93.6 | SRR1805707 |
| | | *Acartia tonsa* | 14 873 | 90.5 | SRR6038073 |
| | **Calanidae** | *Calanoides acutus* | 98 694 | 96.6 | SRR15600794 |
| | | *Calanoides carinatus* | 78 900 | 95.7 | ERR15742263 |
| | | *Calanus finmarchicus* | 92 719 | 95.5 | SRR14700043 |
| | | *Calanus glacialis* | 85 026 | 96.9 | SRR17240410 |
| | | *Calanus helgolandicus* | 28 005 | 93.4 | SRR20015053 |
| | | *Calanus hyperboreus* | 57 524 | 83.2 + 9 | ERR15761856s |
| | | *Calanus marshallae* | 74 237 | 93.4 | SRR17248905 |
| | | *Calanus pacificus* | 120 425 | 96.4 | SRR17249508 |
| | | *Calanus propinquus* | 110 660 | 85.6 | SRR12855516 |
| | | *Calanus sinicus* | 48 251 | 90.3 | SRR1023631 |
| | | *Neocalanus cristatus* | 50 832 | 91.9 | SRR12968742 |
| | | *Neocalanus flemingeri* | 69 883 | 94.4 | SRR5873560 |
| | | *Neocalanus plumchrus* | 42 539 | 92.1 | SRR13083233 |
| | ***Clausocalanidae*** | *Pseudocalanus sp.* | 47 129 | 88.2 + 5.2 | ERR15761855s |
| | **Diaptomidae** | *Gigantodiaptomus amblyodon* | 15 579 | 81.9 | SRR1653196 |
| | **Eucalanidae** | *Eucalanus hyalinus* | 43 025 | 91.7 | ERR15742256 |
| | | *Eucalanus bungi* | 22 032 | 69,5+14,1 | SRR13648609 |
| | **Rhincalanidae** | *Rhincalanus gigas* | 77 312 | 95.9 | SRR12732908 |
| | **Euchaetidae** | *Euchaeta spp.* | 133 666 | 96.5 | SRR13648606 |
| | **Metridinidae** | *Metridia longa* | 47 860 | 93.9 | ERR15761857s |
| | | *Metridia lucens* | 54 766 | 90 | SRR12968740 |
| | | *Metridia pacifica* | 54 955 | 94.3 | SRR12968740 |
| | | *Pleuromamma abdominalis* | 141 046 | 96.1 | SRR13648607 |

| | | | | | |
|---|---|---|---|---|---|
| | | *Pleuromamma robusta* | 44 973 | 91.8 | ERR15742262 |
| | | *Pleuromamma xiphias* | 135 093 | 82.9 +13.3 | SRR5062200 |
| | **Pontellidae** | *Labidocera madurae* | 43 768 | 95.5 | SRR5922844 |
| | **Pseudodiaptimidae** | *Pseudodiaptomus annandalei* | 39 139 | 88.2 + 5.2 | SRR4436675 |
| | **Temoridae** | *Eurytemora affinis* | 26 575 | 96 | SRR2551713 |
| | | *Eurytemora carolleeae* | 24 170 | 96.4 | SRR2551712 |
| | | *Temora longicornis* | 26 001 | 92.7 | SRR20015055 |
| | | *Temora stylifera* | 52 695 | 88.8 + 7.6 | SRR11788221 |
| Cyclopoida | **Cyclopettidae** | *Paracyclopina nana* | 22 981 | 97.9 | SRR1664735 |
| | **Cyclopidae** | *Apocyclops royi* | 20 944 | 57.7 + 19.8 | ERR2811089 |
| | | *Eucyclops serrulatus* | 22 760 | 87.9 + 4.8 | SRR1050725 |
| | **Lernaidea** | *Lernaea cyprinacea* | 77 311 | 91.6 | SRR1107498 |
| | **Oithonidae** | *Oithona nana* | 23 141 | 95.3 | ERR3503858 |
| | | *Oithona similis* | 23 139 | 95.1 | ERR3255855 |
| Harpacticoida | **Harpacticidae** | *Tigriopus californicus* | 29 825 | 96.6 | SRR1613952 |
| | | *Tigriopus japonicus* | 24 055 | 98.6 | SRR1784985 |
| | | *Tigriopus kingsejongensis* | 23 856 | 96 | SRR2034718 |
| | **Laophontidae** | *Platychelipus littoralis* | 99 632 | 97 | SRR10208652 |
| | **Tisbidae** | *Tisbe furcata* | 26 018 | 67.4+19.2 | ERR2209857 |
| Siphonostomatoida | **Caligidae** | *Caligus clemensi* | 22 529 | 81.8 + 11.2 | SRR13648605 |
| | | *Caligus rogercresseyi* | 34 700 | 93.3 | GAZX01 |
| | | *Caligus confusus* | 22 249 | 95.3 | SRR25611632 |
| | | *Lepeophtheirus salmonis* | 17 637 | 87.5 + 5.3 | HACA01 |
| | **Lerneaopodiae** | *Tracheliastes polycolpus* | 14 634 | 95.2 | GGQW01 |

### Supplementary Table 2. Table of three key functional traits across copepod species and related studies.

Information on three functional traits: pelagic layer, post-embryonic dormancy (noted diapause here), and embryonic dormancy (noted resting eggs here) was compiled for copepod species based on published literature. Symbols indicate the following: “?” denotes that the trait is described in another species of the same genus; “NA” indicates that no information was available in the literature.

| Order | Familly | Organism | Pelagic layer | Layer references | Diapause | Diapause reference | Resting egg | Resting egg reference |
|---|---|---|---|---|---|---|---|---|
| | | *Acartia clausi* | Epipelagic | [F5 ; F6 ; F7] | N | [G1 ; G4 ; G10] | Y | [H1 ; H3] |
| | Acartiidae | *Acartia fossae* | Epipelagic ? | | N | | Y ? | |
| | | *Acartia tonsa* | Epipelagic | [F25] | N | | Y | [H3] |
| | | *Calanoides acutus* | Epimesopelagic | [F6] | Y | [G7] | NA | |
| | | *Calanoides carinatus* | Epibathypelagic | [F5 ; F6] | Y | [G4] | NA | |
| | | *Calanus finmarchicus* | Epimesopelagic | [F20] | Y | [G1 ; G4] | N | [H1] |
| | | *Calanus glacialis* | Epimesopelagic | [F12] | Y | [G1 ; G4] | N | [H1] |
| | | *Calanus helgolandicus* | Epibathypelagic | [F1 ; F2 ; F6] | Y | [G1 ; G4] | N | [H1] |
| | | *Calanus hyperboreus* | Epibathypelagic | [F6 ; F12] | Y | [G1 ; G4] | N ? | |
| | Calanidae | *Calanus marshallae* | Epimesopelagic ? | | Y | [G1 ; G4] | N | [H1] |
| | | *Calanus pacificus* | Epimesopelagic | [F16] | Y | [G1 ; G4] | N | [H1] |
| | | *Calanus propinquus* | Epimesopelagic ? | [F6] | Y | [G10] | N | [H1] |
| | | *Calanus sinicus* | Epimesopelagic | [F24] | N | [G1] | N | [H1] |
| | | *Neocalanus cristatus* | Epibathypelagic | [F6 ; F19] | Y | [G1 ; G4] | N | [H1] |
| | | *Neocalanus flemingeri* | Epimesopelagic ? | [F6] | Y | [G4] | N ? | |
| | | *Neocalanus plumchrus* | Epibathypelagic | [F6 ; F19] | Y | [G4] | N ? | |
| Calanoida | Clausocalanidae | *Pseudocalanus sp.* | Epimesopelagic ? | [F6 ; F22] | N ?? | [G1 ; G4 ; G10] | N ? | [H1] |
| | Diaptomidae | *Gigantodiaptomus amblyodo* | Temporary pond | [F10] | NA | | Y | [H5] |
| | Eucalanidae | *Eucalanus hyalinus* | Epibathypelagic | [F2 ; F6] | Y ? | | NA | |
| | | *Eucalanus bungii* | Epibathypelagic | [F6] | Y | [G4] | NA | |
| | Rhincalanidae | *Rhincalanus gigas* | Epibathypelagic | [F6 ; F15] | Y | [G4] | NA | |
| | Euchaetidae | *Euchaeta spp.* | Epibathypelagic ? | [F21] | N ? | [G10] | NA | |
| | | *Metridia longa* | Epibathypelagic | [F6; F13] | Y | [G4 ; G10] | N | [H1] |
| | | *Metridia lucens* | Epibathypelagic | [F6] | Y | [G4 ; G10] | N ? | |
| | Metridinidae | *Metridia pacifica* | Epibathypelagic | [F6] | Y | [G1 ; G4] | N | [H1] |
| | | *Pleuromamma abdominalis* | Epibathypelagic | [F6 ; F21] | N | [G10] | NA | |
| | | *Pleuromamma robusta* | Epibathypelagic | [F25] | N | [G10] | NA | |
| | | *Pleuromamma xiphias* | Epibathypelagic | [F5 ; F6] | N | [G10] | NA | |
| | Pontellidae | *Labidocera madurae* | Epipelagic | [F14] | NA | | Y | [H3 ; H4] |
| | Pseudodiaptimidae | *Pseudodiaptomus annandale* | Epipelagic | [F6] | N ? | [G1] | NA | |
| | | *Eurytemora affinis* | Epipelagic | [F17] | N ? | [G1] | Y | [H3] |
| | Temoridae | *Eurytemora carolleeae* | Epipelagic ? | | N ? | | NA | |
| | | *Temora longicornis* | Epipelagic | [F6] | N | [G1] | Y | [H1 ; H3] |
| | | *Temora stylifera* | Epipelagic | [F6] | N | [G10] | Y ? | |
| | Cyclopettidae | *Paracyclopina nana* | Epipelagic | | N | [G5] | N | [H4] |
| | Cyclopidae | *Apocyclops royi* | Benthic | [F11] | N | [G5] | N | [H4] |
| | | *Eucyclops serrulatus* | Benthic | [F9] | N ?? | [G5 ; G6] | N | [H4] |
| Cyclopoida | Lernaidea | *Lernaea cyprinacea* | Host | | N | [G5] | N | [H4] |
| | Oithonidae | *Oithona nana* | Epipelagic | [F3 ; F4 ; F6; F7] | N | [G4 ; G5] | N | [H1 ; H4] |
| | | *Oithona similis* | Epipelagic | [F6] | N | [G4 ; G5] | N | [H1 ; H4] |
| | | *Tigriopus californicus* | Benthic | [F11] | N | [G5] | N | [H4] |
| | Harpacticidae | *Tigriopus japonicus* | Benthic | [F11] | N | [G5] | N | [H4] |
| | | *Tigriopus kingsejongensis* | Benthic | [F11] | N | [G5] | N | [H4] |
| Harpacticoida | Laophontidae | *Platychelipus littoralis* | Benthic | [F11] | N | [G5] | N | [H4] |
| | Canthocamptidae | *Cletocamptus dominicanus* | Benthic | | N ? | [G9] | NA | |
| | Tisbidae | *Tisbe furcata* | Benthic | | NA | | NA | |
| | | *Tisbe holothuriae* | Benthic | [F11] | NA | | NA | |
| | | *Caligus clemensi* | Host | | N | [G5] | N | [H4] |
| | Caligidae | *Caligus rogercresseyi* | Host | | N | [G5] | N | [H4] |
| Siphonostomatoida | | *Caligus confusus* | Host | | N ? | | N | [H4] |
| | | *Lepeophtheirus salmonis* | Host | | N | [G5] | N | [H4] |
| | Lerneaopodiae | *Tracheliastes polycolpus* | Host | | N | [G5] | N | [H4]+D2:I53 |

#### Supplementary Table 3. Table of references of the three key functional traits

| F | | Layer references |
|---|---|---|
| F | 1 | Andersen V, Gubanova A, Nival P, Ruellet T. Zooplankton community during the transition from spring bloom to oligotrophy in the open NW Mediterranean and effects of wind events. 2. Vertical distributions and migrations. Journal of Plankton Research. 2001;23(3):243-61. |
| F | 2 | Andersen V, Devey C, Gubanova A, Picheral M, Melnikov V, Tsarin S, et al. Vertical distributions of zooplankton across the Almeria–Oran frontal zone (Mediterranean Sea). Journal of Plankton Research. 2004;26(3):275-93. |
| F | 3 | Böttger-Schnack R. Community structure and vertical distribution of cyclopoid copepods in the Red Sea. Marine Biology. 1990;106(3):473-85. |
| F | 4 | Böttger-Schnack R. Vertical structure of small metazoan plankton, especially noncalanoid copepods. I. Deep Arabian Sea. Journal of Plankton Research. 1996;18(7):1073-101. |
| F | 5 | Boxshall GA, Halsey SH. An introduction to copepod diversity: Ray Society; 2004. |
| F | 6 | Razouls C., Desreumaux N., Kouwenberg J. and de Bovée F., 2005-2023. - Biodiversity of Marine Planktonic Copepods (morphology, geographical distribution and biological data). Sorbonne University, CNRS. Available at http://copepodes.obs-banyuls.fr/en |
| F | 7 | Scotto di Carlo B, Ianora A, Fresi E, Hure J. Vertical zonation patterns for Mediterranean copepods from the surface to 3000 m at a fixed station in the Tyrrhenian Sea. Journal of Plankton Research. 1984;6(6):1031-56. |
| F | 8 | Atkinson, Angus, P. Ward, Adrian Hill, Andrew Brierley, and GC Cripps. 'Krill-Copepod Interactions at South Georgia, Antarctica, II. Euphausia Superba as a Major Control on Copepod Abundance'. Marine Ecology-Progress Series - MAR ECOL-PROGR SER 176 (18 January 1999): 63–79. https://doi.org/10.3354/meps176063. |
| F | 9 | Di Lorenzo, Tiziana, Walter di marzio, Marco Cifoni, Barbara Fiasca, Mariella Baratti, Maria Sáenz, and Diana Galassi. 'Temperature Effect on the Sensitivity of the Copepod Eucyclops Serrulatus (Crustacea, Copepoda, Cyclopoida) to Agricultural Pollutants in the Hyporheic Zone'. Current Zoology 61 (1 August 2015): 629–40. https://doi.org/10.1093/czoolo/61.4.629. |
| F | 10 | Dussart, Bernard. 'Les Copépodes Des Eaux Continentales d'Europe Occidentale. Tome I: Calanoides et Harpacticoides. 210 Fig. Paris: Editions N. Boublée & Cie, 1967. 500 Pp. 108. |
| F | 11 | Encyclopedia of Life. Available from http://eol.org. Accessed Feb 2023. |
| F | 12 | Falk-Petersen, Stig, Eva Leu, Jørgen Berge, Slawomir Kwasniewski, Henrik Nygård, Anders Røstad, Essi Keskinen, et al. 'Vertical Migration in High Arctic Waters during Autumn 2004'. Deep Sea Research Part II: Topical Studies in Oceanography, Carbon flux and ecosystem feedback in the northern Barents Sea in an era of climate change, 55, no. 20 (1 October 2008): 2275–84. https://doi.org/10.1016/j.dsr2.2008.05.010. |
| F | 13 | Geinrikh, A.K., K. N. Kosobokova, and Yu. A. Rudyakov. 'Seasonal variations in the vertical distribution of some prolific copepods of the Arctic basin'. Biol. Tsentral'nogo Arkticheskogo Basseina, 1980. |
| F | 14 | Hassett, R. P., and G. W. Boehlert. 'Spatial and Temporal Distributions of Copepods to Leeward and Windward of Oahu, |

Hawaiian Archipelago'. Marine Biology 134, no. 3 (1 August 1999): 571–84. https://doi.org/10.1007/s002270050572.
Hopkins, Thomas L., Thomas M. Lancraft, Joseph J. Torres, and Joseph Donnelly. 'Community Structure and Trophic Ecology of Zooplankton in the Scotia Sea Marginal Ice Zone in Winter (1988)'. Deep Sea Research Part I: Oceanographic Research Papers 40, no. 1 (1 January 1993): 81–105.
F 15 https://doi.org/10.1016/0967-0637(93)90054-7.
Johnson, Catherine Lynn, and David M. Checkley. 'Vertical Distribution of Diapausing Calanus Pacificus (Copepoda) and Implications for Transport in the California Undercurrent'. Progress in Oceanography 62, no. 1 (1 July 2004): 1–13.
F 16 https://doi.org/10.1016/j.pocean.2004.08.001.
Labuce, Astra, Solvita Strake, and Inta Dimante-Deimantovica. 'Eurytemora Affinis (Poppe, 1880) (Copepoda, Calanoida) in the Gulf of Riga, Baltic Sea — Elemental Composition and Diurnal Vertical Migration'. Crustaceana 93, no. 3–5 (8 June 2020):
F 17 447–66. https://doi.org/10.1163/15685403-00003977.
Lo, Wen-Tseng, Chang-Tai Shih, and Jiang-Shiou Hwang. 'Diel Vertical Migration of the Planktonic Copepods at an Upwelling Station North of Taiwan, Western North Pacific'. Journal of Plankton Research 26, no. 1 (1 January 2004): 89–97.
F 18 https://doi.org/10.1093/plankt/fbh004.
Miller, Charles B., Bruce W. Frost, Harold P. Batchelder, Martha J. Clemons, and Richard E. Conway. 'Life Histories of Large, Grazing Copepods in a Subarctic Ocean Gyre: Neocalanus Plumchrus, Neocalanus Cristatus, and Eucalanus Bungii in the Northeast Pacific'. Progress in Oceanography 13, no. 2 (January 1984):
F 19 201–43. https://doi.org/10.1016/0079-6611(84)90009-0.
Ohman, Mark D., and Jefrey A. Runge. 'Sustained Fecundity When Phytoplankton Resources Are in Short Supply: Omnivory by Calanus Finmarchicus in the Gulf of St. Lawrence'. Limnology and Oceanography 39, no. 1 (1994): 21–36.
F 20 https://doi.org/10.4319/lo.1994.39.1.0021.
Takenaka, Yasuhiro, Atsushi Yamaguchi, Naoki Tsuruoka, Masaki Torimura, Takashi Gojobori, and Yasushi Shigeri. 'Evolution of Bioluminescence in Marine Planktonic Copepods'. Molecular Biology and Evolution 29, no. 6 (1 June 2012): 1669–81.
F 21 https://doi.org/10.1093/molbev/mss009.
Turner, Jefferson T., and Michael J. Dagg. 'Vertical Distributions of Continental Shelf Zooplankton in Stratified and Isothermal Waters'. Biological Oceanography 3, no. 1 (1 January 1983): 1–40.
F 22 https://doi.org/10.1080/01965581.1983.10749470.
WoRMS Editorial Board (2023). World Register of Marine Species. Available from https://www.marinespecies.org at VLIZ. Accessed
F 23 2023-02. doi:10.14284/170
Yoshida, T., T. Toda, V. Kuwahara, S. Taguchi, and B. H. R. Othman. 'Rapid Response to Changing Light Environments of the Calanoid Copepod Calanus Sinicus'. Marine Biology 145, no. 3 (1 September 2004): 505–13.
F 24 https://doi.org/10.1007/s00227-004-1343-5.
Pata, P.R. and Hunt, B.P.V. (2024), Harmonizing marine zooplankton trait data toward a mechanistic understanding of ecosystem functioning. Limnol Oceanogr.
F 25 https://doi.org/10.1002/lno.12478

#### G Post-embryonic dormancy reference

Barton, Andrew, A. Pershing, Elena Litchman, Nicholas Record, Kyle Edwards, Zoe Finkel, Thomas Kiørboe, and Ben Ward. 'The Biogeography of Marine Plankton Traits'. Ecology Letters 16 (30
G 1 January 2013). https://doi.org/10.1111/ele.12063.
Baumgartner, Mark F., and Ann M. Tarrant. 'The Physiology and Ecology of Diapause in Marine Copepods'. Annual Review of Marine Science 9, no. 1 (2017): 387–411.
G 2 https://doi.org/10.1146/annurev-marine-010816-060505.
Bradford-Grieve, Janet M., and Shane T. Ahyong. 'Phylogenetic Relationships among Genera in the Calanidae (Crustacea:
G 3 Copepoda) Based on Morphology'. Journal of Natural History 44,

| | | |
|---|---|---|
| | | no. 5–6 (22 February 2010): 279–99. https://doi.org/10.1080/00222930903359644. |
| G | 4 | Brun, Philipp, Mark R. Payne, and Thomas Kiørboe. 'A Trait Database for Marine Copepods'. Supplement to: Brun, P et al. (2017): A Trait Database for Marine Copepods. Earth System Science Data, 9(1), 99-113, Https://Doi.Org/10.5194/Essd-9-99-2017. PANGAEA, 11 July 2016. https://doi.org/10.1594/PANGAEA.862968. |
| G | 5 | Dahms, Hans-Uwe. 'Dormancy in the Copepoda — an Overview'. Hydrobiologia 306 (1 June 1995): 199–211. https://doi.org/10.1007/BF00017691. |
| G | 6 | Maier, Gerhard. 'The Effect of Temperature on the Development, Reproduction, and Longevity of Two Common Cyclopoid Copepods — Eucyclops Serrulatus (Fischer) and Cyclops Strenuus (Fischer)'. Hydrobiologia 203, no. 3 (1 September 1990): 165–75. https://doi.org/10.1007/BF00005685. |
| G | 7 | Pinti, J., Jónasdóttir, S.H., Record, N.R. and Visser, A.W. (2023), The global contribution of seasonally migrating copepods to the biological carbon pump. Limnol Oceanogr, 68: 1147-1160. https://doi.org/10.1002/lno.12335 |
| G | 8 | 'Copepod (Crustacea) Emergence from Soils from Everglades Marshes with Different Hydroperiods'. Accessed 2 October 2023. https://www.tandfonline.com/doi/epdf/10.1080/02705060.2000.9663774?needAccess=true. |
| G | 9 | Loftus, William F., and Janet W. Reid. 'Copepod (Crustacea) Emergence from Soils from Everglades Marshes with Different Hydroperiods'. Journal of Freshwater Ecology 15, no. 4 (1 December 2000): 515–23. https://doi.org/10.1080/02705060.2000.9663774. |
| G | 10 | McGinty, N., Barton, A. D., Record, N. R., Finkel, Z. V., & Irwin, A. J. (2018). Traits structure copepod niches in the North Atlantic and Southern Ocean. Marine Ecology Progress Series, 601, 109–126. https://www.jstor.org/stable/26503145 |
| **H** | | **Resting egg reference** |
| H | 1 | Barton, Andrew, A. Pershing, Elena Litchman, Nicholas Record, Kyle Edwards, Zoe Finkel, Thomas Kiørboe, and Ben Ward. 'The Biogeography of Marine Plankton Traits'. Ecology Letters 16 (30 January 2013). https://doi.org/10.1111/ele.12063. |
| H | 2 | Baumgartner, Mark F., and Ann M. Tarrant. 'The Physiology and Ecology of Diapause in Marine Copepods'. Annual Review of Marine Science 9, no. 1 (2017): 387–411. https://doi.org/10.1146/annurev-marine-010816-060505. |
| H | 3 | Brun, Philipp, Mark R. Payne, and Thomas Kiørboe. 'A Trait Database for Marine Copepods'. Supplement to: Brun, P et al. (2017): A Trait Database for Marine Copepods. Earth System Science Data, 9(1), 99-113, Https://Doi.Org/10.5194/Essd-9-99-2017. PANGAEA, 11 July 2016. https://doi.org/10.1594/PANGAEA.862968. |
| H | 4 | Dahms, Hans-Uwe. 'Dormancy in the Copepoda — an Overview'. Hydrobiologia 306 (1 June 1995): 199–211. https://doi.org/10.1007/BF00017691. |
| H | 5 | Marrone, Federico, Sabrina Lo Brutto, and Marco Arculeo. 'Molecular Evidence for the Presence of Cryptic Evolutionary Lineages in the Freshwater Copepod Genus Hemidiaptomus G.O. Sars, 1903 (Calanoida, Diaptomidae)'. Hydrobiologia 644, no. 1 (1 May 2010): 115–25. https://doi.org/10.1007/s10750-010-0101-6. |

#### Supplementary Table 4: Ancestral trait reconstruction

The evolutionary history of each trait was reconstructed using the presence–absence matrix and the phylogenetic topology shown in Figure 1. Three comparative methods were applied to estimate the probability of trait acquisition (post-embryonic dormancy, resting eggs, and mesopelagic adaptation), all using default parameters: maximum likelihood under the Mk1 model in *Mesquite*, maximum likelihood under the Asymmetrical Mk model in *Mesquite*, and a Bayesian inference under the MultiState model implemented in *BayesTraits* v5.0.1 (Pagel et al., 2004; Meade and Pagel, 2022).

| Software | Method | Traits | p1(calanoida) | p1(centro\|pan) | p1(centro) | p1(pan) |
|---|---|---|---|---|---|---|
| Mesquite | mk1 | Diapause | 0.863 | 0.822 | 0.139 | 0.97 |
| | | Mesopelagic | 0.56 | 0.556 | 0.008 | 0.997 |
| | | RestingEgg | 0.003 | 0.021 | 0.883 | 0.001 |
| | AsymmMk | Diapause | 0.97 | 0.955 | 0.245 | 0.998 |
| | | Mesopelagic | 0.788 | 0.784 | 0.027 | 0.999 |
| | | RestingEgg | 0 | 0.007 | 0.977 | 0 |
| BayesTraits | MCMC MultiState | Diapause | 0.939 | 0.811 | 0.163 | 0.992 |
| | | Mesopelagic | 0.95 | 0.753 | 0.023 | 0.999 |
| | | RestingEgg | 0.371 | 0.677 | 0.989 | 0.079 |

#### Supplementary Table 5. Classification of upregulated genes

https://github.com/madoui/Hypoxia/blob/main/DE/Supplementary_Table_S4.xlsx

#### Supplementary Table 6. Enzyme symbols

Symbols of enzymes used in Figure 4

| Enzyme | Symbol |
|---|---|
| oxoglutarate deshydrogenase complex subunit 1 | OGCDE1 |
| isocitrate lyase | ICL |
| glutamate deshydrogenase | GLDH |
| glutamine synthetase | GS |
| cysteine dioxygenase | CDO |
| Cysteine sulfinic acid decarboxylase | CSAD |
| sulfoacetaldehyde acetyltransferase | Xsc |
| 2-amino-3-ketobutyrate coenzyme A ligase | KBL |
| Acetoacetyl-CoA synthetase | AACS |
| cystathionine gamma-lyase | CTH |
| Adenosylmethionine decarboxylase | AdoMetDC |
| Glycine N-methyltransferase | GNMT |
| S-Adenosylmethionine synthetase | MAT |
| alpha-aminoadipic semialdehyde synthase, | AASS |
| 3-amino-2-methylpropionate transaminase | AMT |
| 3-amino-2-methylpropanoate | AMP |
| Methylmalonate-semialdehyde dehydrogenase | MSDH |
| 2-Methyl-3-oxopropanoic acid | MOXA |
| aminomethyltransferase | AMMT |
| Betaine--homocysteine S-methyltransferase 1 | BHMT |
| Phenylalanine hydroxylase | PAH |
| Hypoxanthine-guanine phosphoribosyltransferase | HGPRT |
| 4-Hydroxyphenylpyruvate dioxygenase | HPPD |
| Glycine cleavage system H protein | H-protein |
| Glycine cleavage system T protein | T-protein |
| Aspartate aminotransferase | AST |
| Aspartate oxydase | Laspo |
| nitric oxide synthase | NOS |
| Hydroxyacid-oxoacid transhydrogenase | HOT |
| Fumarylacetoacetate hydrolase | FAH |
| Fructose-bisphosphate aldolase | ALDO |
| hexokinase | HK |
| alanine deshydrogenase | AD |
| lactate deshydrogenase | LDH |
| (S)-3-amino-2-methylpropionate transaminase | AMPT |
| methionine synthase | MS |
| glutathione synthetase | GSS |

#### Supplementary Table 7: Copepod sampling for newly assembled transcriptomes.

| Species | Date | Latitude | Longitude | Depth | Informations |
|---|---|---|---|---|---|
| Calanoides carinatus | 2010/10/05 | 18° 59.590' S | 10° 19.839' E | 0 - 5 | C5 |
| Eucalanus hyalinus | 2010/10/08 | 23° 00.100' S | 13° 03.304' E | 0-5 | F ; high oxygen |
| Eucalanus hyalinus | 2010/10/08 | 23° 00.100' S | 13° 03.304' E | 250-370 | F ; low oxygen |
| Pleuromamma robusta | 2010/09/27 | 18° 59.590' S | 10° 19.839' E | 200-500 | F |

#### Supplementary Table 8 : Five fossil calibrations for arthropods from Wolfe et al. 2016

| **Node** | **Fossil Species** | **Minimum Age (Ma)** | **Soft Maximum Age (Ma)** |
|---|---|---|---|
| Multicrustacea | *Arenosicaris inflata* | 487 | 636,1 |
| Copepoda/ Siphonostomatoida | *Kabatarina pattersoni* | 93,7 | 521 |
| Copepoda/Podoplea/ Harpaticoida | *Canthocamptidae* | 298 | 521 |
| Thecostraca | *Rhamphoverritor reduncus* | 429,8 | 521 |
| Malacostraca | *Cinerocaris magnifica* | 429,8 | 521 |

#### Supplementary Figure 1: Maximum-likelihood phylogenetic tree of isocitrate lyase.

Genes highlighted in blue are from bacterial origin

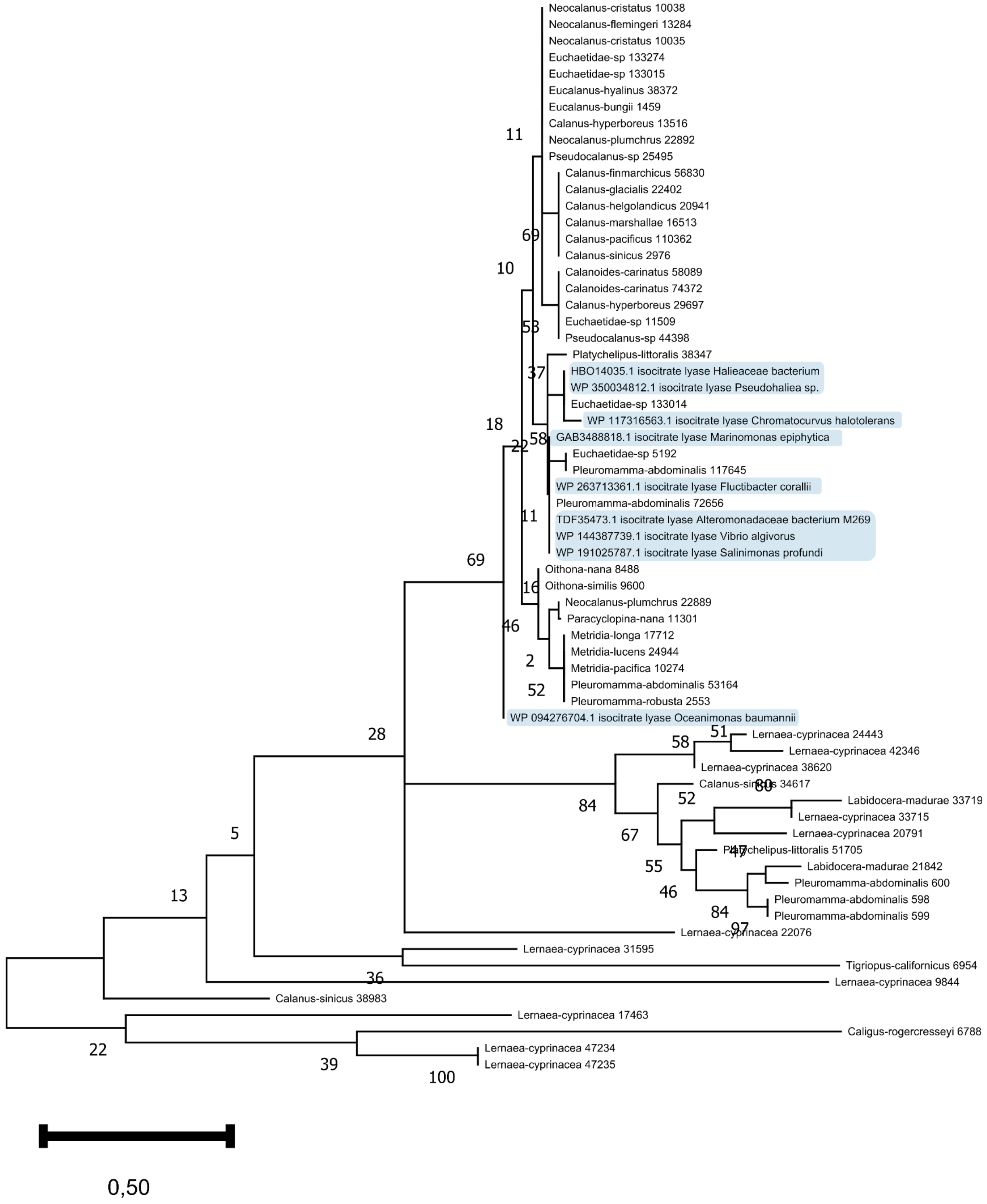

Supplementary Figure 2: MA plot of *Eucalanus hyalinus* transcripts

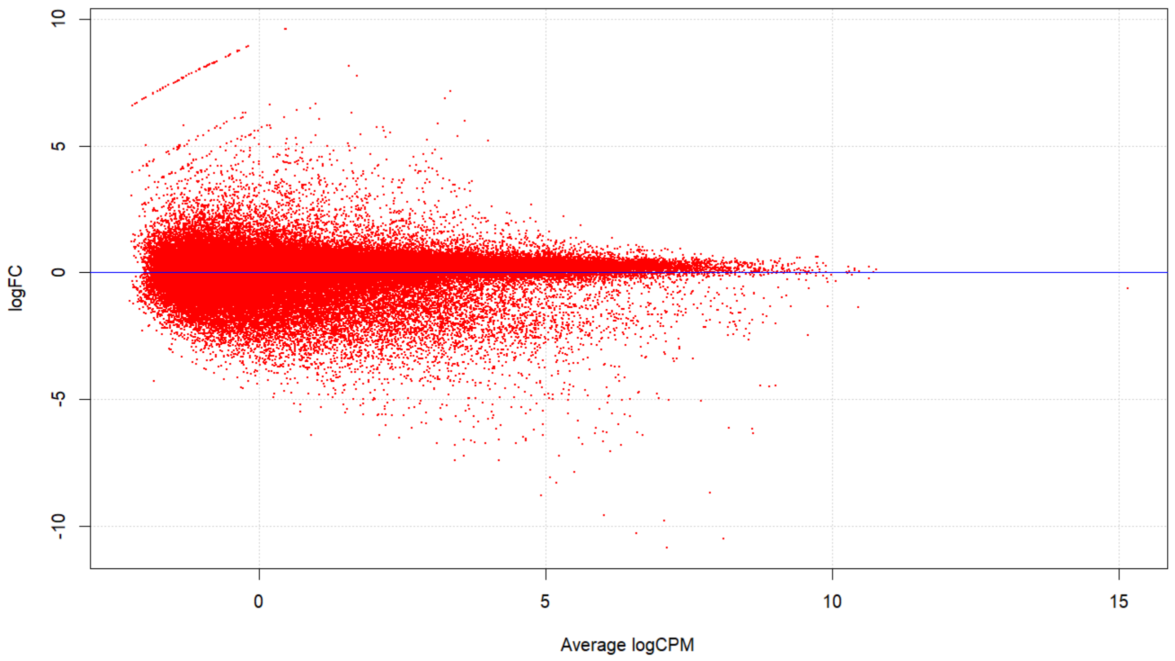

### Supplementary Figure 3: Convergence analysis between the three independent MCMC analyses

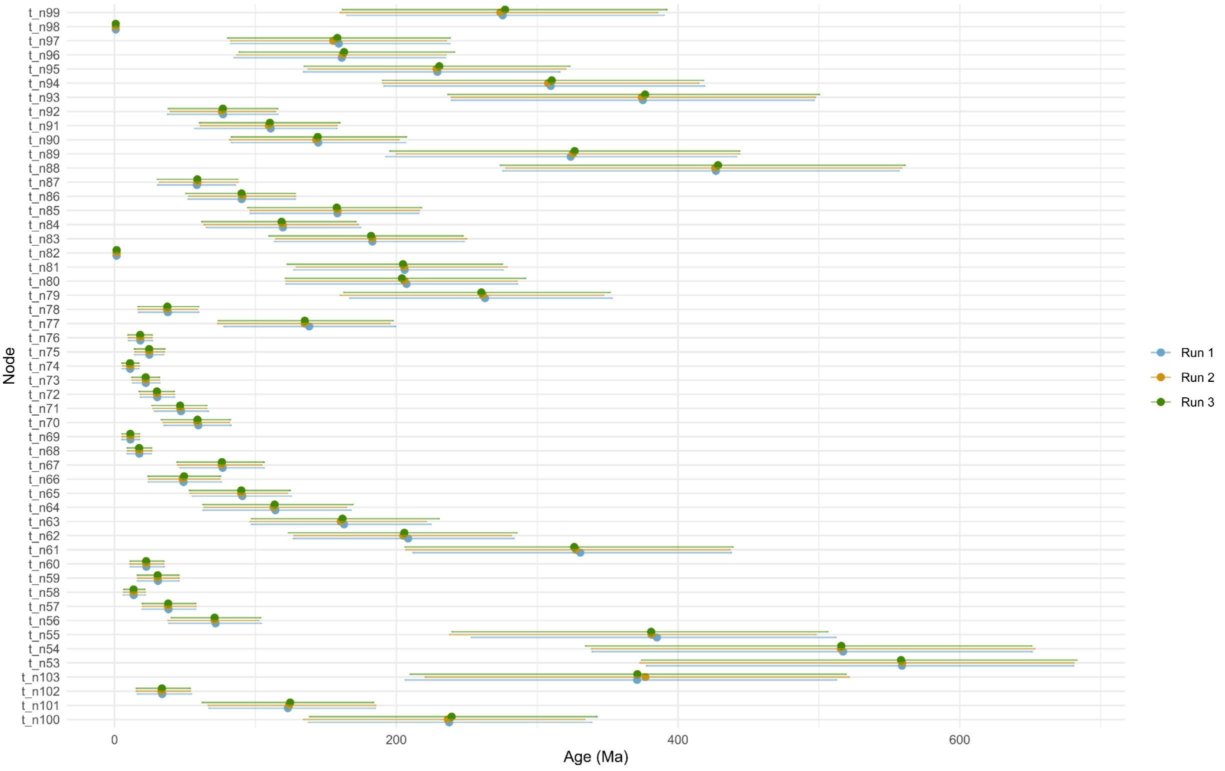

#### Supplementary Figure 4: MCMCTree prior vs posterior highest posterior density widths

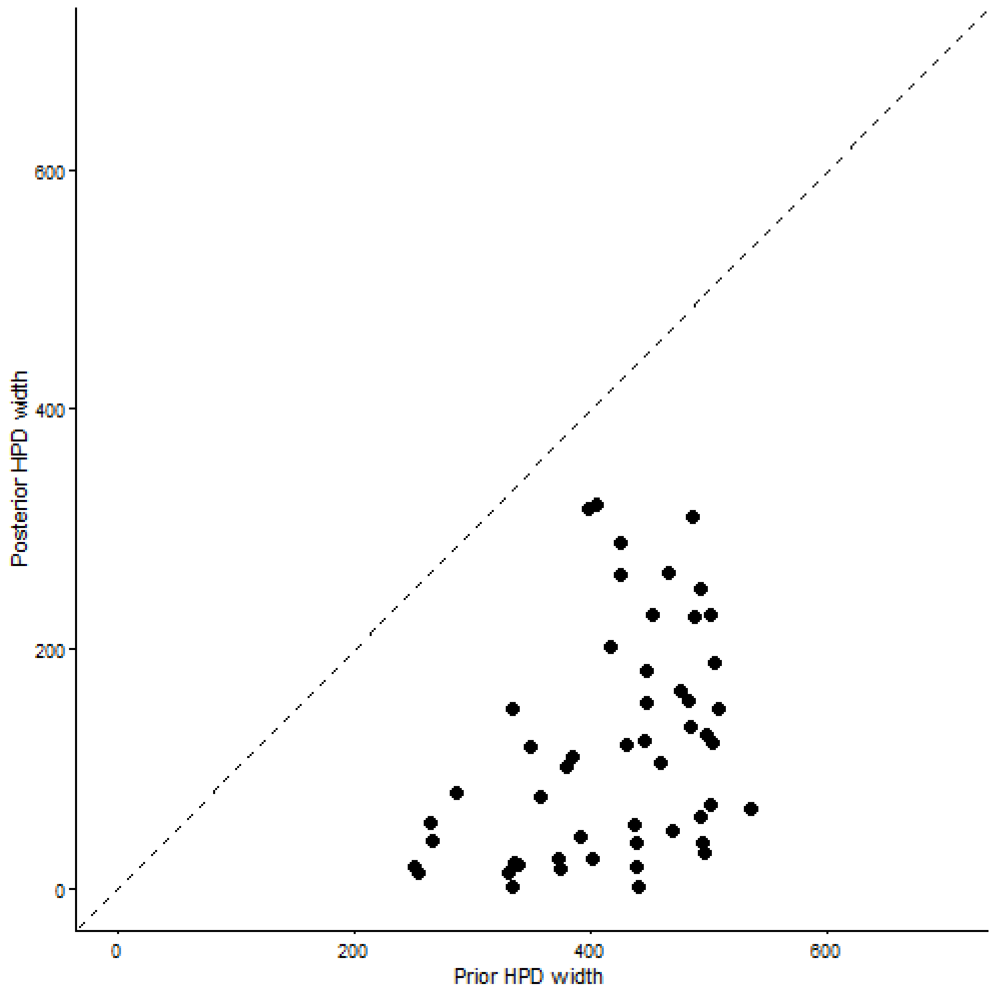

#### Supplementary Results: Transcriptomic response of Eucalanus hyalinus in OMZ

##### Hormonal pathway regulation

We identified the overexpression of an ecdysone kinase, an enzyme that phosphorylates and inactivates ecdysone, the key moulting and metamorphosis hormone in arthropods. In *C. finmarchicus*, the role of ecdysone signalling during diapause has been demonstrated through downregulation of the ecdysone receptor[1]. We also observed upregulation of a juvenile hormone–binding protein (JHBP) and a methyl farnesoate (MF) epoxidase. While the presence of juvenile hormone (JH) in crustaceans remains uncertain, its precursor MF is widespread in decapods and controls vitellogenesis and ovarian maturation[2], and inactivation of MF by JHBP has been demonstrated in crabs[3]. Together, these three enzymes appear to block hormonal signals for moulting and reproduction, suggesting developmental arrest in OMZ with developmental arrest of the CV juveniles. A *sleepless/quiver* ortholog was also overexpressed. Quiver is known to decrease neuronal excitability in Drosophila[4], which suggests a neurophysiological mechanism for reducing metabolic demand during hypoxia, through reduction of stimulus-induced motility, but functional validation is required to confirm this hypothesis.

##### Extracellular matrix remodelling

In addition, 12 genes encoding secreted proteases predicted to target the extracellular matrix (ECM), along with three fibrillar collagen genes and one fibronectin, were up-regulated in OMZ. ECM remodelling is a hallmark of hypoxia responses and is conserved in both mammals[5] and flatworms[6] suggesting that it is also induced by hypoxia in *E. hyalinus*. We identified 27 proteins involved in the transport of lipid (lipocalin), sugar, amino acid, ion and iron (ferritin and ferroportin)

(Supplementary Table 5), which supports a reallocation of released amino acids and other metabolites from the ECM through active transport and cellular uptake.

The ECM glycoaminoglycans are remodelled by sulfation modification through the activation of sulfotransferase and arylsulfatase (Fig. 4c). Similar sulfotransferase activity has already been observed in the deep-sea shrimp[7,8] and barnacle[9] species, which illustrates evolutionary convergence of gene duplication in hypoxia-tolerant marine crustaceans. An outcome of this pathway may be the generation of reduced sulphur compounds, particularly hydrogen sulphide ($H_2S$), through microbial sulphate reduction. $H_2S$ is a known inhibitor of cytochrome c oxidase and has been implicated in the induction of metabolic torpor in overwintering vertebrates[10,11]. Sulphate-reducing bacteria (SRB), capable of producing $H_2S$, have been found in high abundance in the gut microbiota of overwintering insects[12] and of the mesopelagic copepod *Pleuromamma* spp.[13], raising the possibility that SRB-derived sulphide contributes to host hypoxia adaptation through metabolic suppression.

**Carbohydrate and lipid metabolic reprogramming**

Two enzymes of the ascorbic acid biosynthetic pathway, including L-gulonolactone oxidase (GULO), were significantly up-regulated under hypoxia (Fig. 4d). Seven lipases involved in the degradation of triacylglycerols and diacylglycerols are up-regulated, releasing fatty acids and glycerol (Fig. 3e). These fatty acids are either reduced into fatty alcohols for wax esters synthesis, or alternatively elongated and desaturated to recycle membrane lipids. This pathway enables the conversion of triacylglycerols into wax esters, which is a hallmark of pre-diapause lipids in Calanidae that use wax esters directly as the primary long-term energy store[14]. Importantly, the transition to wax esters also provides a biophysical mechanism for buoyancy regulation at depth, as shown in *Calanoides acutus*, where pressure-induced phase transitions of wax esters contribute to buoyancy during diapause[15]. Glycolysis is activated through canonical HIF-1 targets including two hexokinases and two aldolases, accompanied by increased lactate production.

**Amino acid metabolic reprogramming**

The amino acid metabolism indicates anaplerotic input into the tricarboxylic acid (TCA) cycle via catabolism of branched-chain, aromatic amino acids and aspartate (Fig. 4f, Supplementary Table 6). The overexpression of an isocitrate lyase (ICL) gene suggests the activation of the glyoxylate shunt leading to gluconeogenesis, and this activation is a hallmark of starvation in *C. elegans*. The phylogeny of ICL from the orthogroup with additional best hit protein sequences suggests a gene acquisition by copepod ancestors via bacterial horizontal gene transfer (HGT) (Supplementary Fig. 1), which is consistent with previous identification of multiple HGTs of the glyoxylate cycle in Metazoa[16]. The pentose phosphate pathway is activated by the overexpression of glucose-1-dehydrogenase, leading to the production of gluconate and the phosphoribosylpyrophosphate (PRPP) synthase overexpression, promoting the conversion of glucose into PRPP. The pentose phosphate pathway has been previously described as activated during diapause preparation in *C. finmarchicus*[17]. This intermediate is essential for nucleotide biosynthesis to support DNA/RNA synthesis and repair processes.

**Antioxydant production**

Up-regulation of the glycine cleavage system fuels the folate and methionine cycles, thereby sustaining cysteine biosynthesis via the transsulfuration pathway. This cysteine supply is critical for the production of multiple antioxidant defences. Among these, glutathione is well documented in copepods as a major thiol redox buffer, with glutathione S-transferase (GST) activity (3 GSTs overexpressed) widely reported under environmental stress[18,19]. Polyamines such as spermidine and spermine play conserved roles in DNA protection and oxidative stress resistance, as shown by QTL analysis in Pacific white shrimp[20]. Sulphur-containing amino acids such as hypotaurine and taurine are known ROS scavengers, membrane stabilisers and maintain glutathione stores[21], with studies in bivalves[22] and shrimps[23] demonstrating protective functions against oxidative stress during hypoxia. Arginine is metabolised to nitric oxide, a key regulator of hypoxia response that induces muscle relaxation[24] and its production is sustained by the biosynthesis of the essential cofactor tetrahydrobiopterin ($BH_4$). Together, these metabolic rewiring support the active production of at least seven antioxidants to control reactive oxygen (ROS) damage, enhance anaerobic ATP production, and reduce the ATP demand through muscle relaxation.

**Immune system activation**

The up-regulation of sixteen lectin-domain-containing proteins and four Tumor Necrosis Factor (TNF) domain-containing proteins suggests an activation of the innate immune system[25], consistent with different microbial challenges between the epi and the mesopelagic zone[26].

#### References


1. Clark, K. A. J., Brierley, A. S., Pond, D. W. & Smith, V. J. Changes in seasonal expression patterns of ecdysone receptor, retinoid X receptor and an A-type allatostatin in the copepod, *Calanus finmarchicus,* in a sea loch environment: An investigation of possible mediators of diapause. *Gen. Comp. Endocrinol.* **189**, 66–73 (2013).
2. Laufer, H. *et al.* Identification of a Juvenile Hormone-Like Compound in a Crustacean. *Science* **235**, 202–205 (1987).
3. Zhao, M., Wang, W., Ma, C., Zhang, F. & Ma, L. Characterization and expression analysis of seven putative JHBPs in the mud crab *Scylla paramamosain*: Putative relationship with methyl farnesoate. *Comp. Biochem. Physiol. B Biochem. Mol. Biol.* **241**, 110390 (2020).
4. Koh, K. *et al.* Identification of SLEEPLESS, a Sleep-Promoting Factor. *Science* **321**, 372–376 (2008).
5. Higgins, D. F. *et al.* Hypoxia promotes fibrogenesis in vivo via HIF-1 stimulation of epithelial-to-mesenchymal transition. *J. Clin. Invest.* **117**, 3810–3820 (2007).
6. Vozdek, R., Long, Y. & Ma, D. K. The receptor tyrosine kinase HIR-1 coordinates HIF-independent responses to hypoxia and extracellular matrix injury. *Sci. Signal.* **11**, eaat0138 (2018).
7. Zhang, J., Sun, Q., Luan, Z., Lian, C. & Sun, L. Comparative transcriptome analysis of Rimicaris sp. reveals novel molecular features associated with survival in deep-sea hydrothermal vent. *Sci Rep* **7**, 2000 (2017).
8. Zhu, F.-C. *et al.* Insights into the strategy of micro-environmental adaptation: Transcriptomic analysis of two alvinocaridid shrimps at a hydrothermal vent. *PLOS ONE* **15**, e0227587 (2020).
9. Gan, Z. *et al.* Comparative transcriptomic analysis of deep- and shallow-water barnacle species (Cirripedia, Poecilasmatidae) provides insights into deep-sea adaptation of sessile crustaceans. *BMC Genomics* **21**, 240 (2020).
10. Revsbech, I. G. *et al.* Hydrogen sulfide and nitric oxide metabolites in the blood of free-ranging brown bears and their potential roles in hibernation. *Free Radic. Biol. Med.* **73**, 349–357 (2014).

11. Jensen, B. S. *et al.* Suppression of mitochondrial respiration by hydrogen sulfide in hibernating 13-lined ground squirrels. *Free Radic. Biol. Med.* **169**, 181–186 (2021).

12. Wang, J., Chen, H. & Tang, M. Community structure of gut bacteria of Dendroctonus armandi (Coleoptera: Curculionidae: Scolytinae) larvae during overwintering stage. *Sci. Rep.* **7**, 14242 (2017).

13. Sadaiappan, B., PrasannaKumar, C., Nambiar, V. U., Subramanian, M. & Gauns, M. U. Meta-analysis cum machine learning approaches address the structure and biogeochemical potential of marine copepod associated bacteriobiomes. *Sci. Rep.* **11**, 3312 (2021).

14. Berger, C. A., Steinberg, D. K., Copeman, L. A. & Tarrant, A. M. Comparative analysis of the molecular starvation response of Southern Ocean copepods. *Mol. Ecol.* **34**, e17371 (2025).

15. Pond, D. W. The physical properties of lipids and their role in controlling the distribution of zooplankton in the oceans. *J. Plankton Res.* **34**, 443–453 (2012).

16. Kondrashov, F. A., Koonin, E. V., Morgunov, I. G., Finogenova, T. V. & Kondrashova, M. N. Evolution of glyoxylate cycle enzymes in Metazoa: evidence of multiple horizontal transfer events and pseudogene formation. *Biol. Direct* **1**, 31 (2006).

17. Tarrant, A. M. *et al.* Transcriptional Profiling of Metabolic Transitions during Development and Diapause Preparation in the Copepod Calanus finmarchicus. https://doi.org/10.1093/icb/icw060 doi:10.1093/icb/icw060.

18. Lauritano, C., Carotenuto, Y. & Roncalli, V. Glutathione S-Transferases in Marine Copepods. *J. Mar. Sci. Eng.* **9**, 1025 (2021).

19. Roncalli, V. *et al.* Experimental analysis of development, lipid accumulation and gene expression in a high-latitude marine copepod. *J. Plankton Res.* **45**, 885–898 (2023).

20. Li, X. *et al.* Identification of QTLs and candidate genes screening for hypoxia tolerance in Pacific white shrimp (*Litopenaeus vannamei*). *Aquac. Rep.* **42**, 102824 (2025).

21. Baliou, S. *et al.* Protective role of taurine against oxidative stress (Review). *Mol. Med. Rep.* **24**, 1–19 (2021).

22. Haider, F., Falfushynska, H. I., Timm, S. & Sokolova, I. M. Effects of hypoxia and reoxygenation on intermediary metabolite homeostasis of marine bivalves Mytilus edulis and Crassostrea gigas. *Comp. Biochem. Physiol. A. Mol. Integr. Physiol.* **242**, 110657 (2020).
23. Shi, M. *et al.* Effects of taurine supplementation in low fishmeal diet on growth, immunity and intestinal health of *Litopenaeus vannamei*. *Aquac. Rep.* **32**, 101713 (2023).
24. Umbrello, M., Dyson, A., Feelisch, M. & Singer, M. The key role of nitric oxide in hypoxia: hypoxic vasodilation and energy supply-demand matching. *Antioxid. Redox Signal.* **19**, 1690–1710 (2013).
25. Sánchez-Salgado, J. L. *et al.* Participation of lectins in crustacean immune system. *Aquac. Res.* **48**, 4001–4011 (2017).
26. Yu, Z., Yang, J., Liu, L., Zhang, W. & Amalfitano, S. Bacterioplankton community shifts associated with epipelagic and mesopelagic waters in the Southern Ocean. *Sci. Rep.* **5**, 12897 (2015).
